# aniSNA : An R package to assess bias and uncertainty in social networks obtained from animals sampled via direct observations or satellite telemetry

**DOI:** 10.1101/2024.05.10.593659

**Authors:** Prabhleen Kaur, Simone Ciuti, Adele K. Reinking, Jeffrey L. Beck, Michael Salter-Townshend

## Abstract

Animal social network analysis using GPS telemetry datasets provides insights into group dynamics, social structure, and interactions of the animal communities. It aids conservation by characterizing key aspects of animal sociality - including spatially explicit information on where sociality occurs (e.g., habitats, migratory corridors), contributing to informed management strategies for wildlife populations. The aniSNA package provides functions to assess and leverage data collected by sampling a subset of an animal population to perform social network analysis. The methodologies offered in this package are compatible with a variety of location and grouping data, collected through various means (e.g., direct observations, biologgers), however, they are particularly well suited to autocorrelated data streams such as data collected through GPS telemetry radio collars. The techniques assess the data’s suitability to extract reliable statistical inferences from social networks and compute uncertainty estimates around the network metrics in the scenario where a fraction of the population is monitored. The package functions are user-friendly and allow for the implementation of pre-network data permutations for auto-correlated data streams, sensitivity analysis under downsampling, bootstrapping to establish confidence intervals for global and node-level network metrics, and correlation and regression analysis to assess the robustness of node-level network metrics. Using this package, animal ecologists will be able to compute social network metrics, both at the population and individual level, assess their reliability, and use such metrics in further analyses, e.g., to study social network variation within and across populations or link individual sociality to life history. This software also has plotting features that allow for visual interpretation of the findings.

## 1. Introduction: Social network analysis of animal societies

Over the last decade, social network theory has emerged as a competent means to enhance our understanding of complex interactions between animals (Farine and Strandburg-Peshkin 2015a; Hobson, Silk, Fefferman, Larremore, Rombach, Shai, and Pinter-Wollman 2021; Silk, Croft, Delahay, Hodgson, Weber, Boots, and McDonald 2017; Gomes, Boogert, and Cardoso 2023; Torfs, Stevens, Verspeek, Laméris, Guéry, Eens, and Staes 2023; Davis, Crofoot, and Farine 2018; Balasubramaniam, Beisner, Berman, Marco, Duboscq, Koirala, Majolo, MacIntosh, McFarland, Molesti, Ogawa, Petit, Schino, Sosa, Sueur, Thierry, de Waal, and McCowan 2018). Consequently, there has been an upsurge in software and tools available to construct and analyse animal social networks. In `R` (R Core Team 2022), packages have played a crucial role in the analysis of animal social networks by providing a comprehensive set of tools, functions, and methodologies tailored to network analysis. Apart from the general packages available to analyse social networks such as igraph (Csardi and Nepusz 2006), sna (Butts 2020), network (Butts 2015, 2008), statnet(Handcock, Hunter, Butts, Goodreau, Krivitsky, and Morris 2018; Hunter, Handcock, Butts, Goodreau, and Morris 2008), specialised software packages developed to analyse animal networks have surfaced (Farine 2013; Bonnell 2023; Silk, McDonald, Delahay, Padfield, and Hodgson 2020; Sosa, Puga-Gonzalez, Hu, Pansanel, Xie, and Sueur 2020; Ross, McElreath, and Redhead 2023; Silk and Gimenez 2023). Ecologists now have a very solid platform to undertake many types of studies and develop networks from data obtained in various ways owing to this softwares. Most of these software cater to a different aspect of the animal social network studies, such as assistance with importing different data formats and data cleaning, network formation and visualisation, calculation of different network metrics, statistical analysis, community detection, and temporal analysis.

These `R` packages seamlessly integrate with other statistical and data analysis tools in the `R` ecosystem, allowing researchers to combine network analysis with other types of analyses.

The asnipe package (Farine 2013) is one of the first such packages in the `R` environment that provided a novel approach for estimating re-association rates of time between frequently sampled individuals, bridging a gap in the tools available to perform permutation-based statistical testing on animal social network data. The spatsoc (Robitaille, Webber, and Wal 2019) package in `R` facilitates spatial social network analysis using animal telemetry data. It provides flexible functions for generating edge lists, gambit-of-the-group data, data-stream randomization, and group by individual matrices, allowing users of animal telemetry data to generate efficient and intuitive social networks. NetTS (Bonnell 2023) is a time-aggregated network package that uses an adjustable moving window to measure how a social network changes through time. It provides tools for choosing window sizes, comparing observed network measures to null models, and simulating network data to aid in statistical model construction and testing. The package CMRnet (Silk et al. 2020) assists in generating social and movement networks from long-term capture-mark-recapture data, providing insights into demography and behaviour in wild animal populations. Finally, The ANTs (Sosa et al. 2020) package presents itself as the fastest computing environment and an all-in-one toolbox for implementing various social network analysis techniques in use today. The package attempts to manage the limitations of each of its predecessors which are discussed in more detail in Sosa *et al*. (2020). The all-in-one toolbox provides a variety of functions including calculating network formation and network metrics, performing pre-and post-network data randomization, and implementing various statistical tests. Two of the recent `R` packages STRAND (Ross *et al*. 2023) and genNetDem (Silk and Gimenez 2023) in `R` allow to apply generative network models. The STRAND package allows the integration of stochastic block models with social relation models for Bayesian analysis of animal social networks. Package genNetDem simulates integrated network-demographic datasets, generating populations and social networks with known statistical relationships.

### 1.1. Assessing data suitability : samples from a population

By leveraging these features available through different packages, researchers can efficiently analyze complex animal social networks, uncover patterns of interaction, and gain a deeper understanding of the social dynamics within animal populations. The availability of diverse `R` packages has allowed researchers to choose the tools that best suit their specific research questions and study designs. However, one of the concerns before using a dataset to answer a particular research question is the dataset’s adequacy to perform social network analysis and obtain correct inferences (Farine and Strandburg-Peshkin 2015b). Data collection through telemetry devices is rapidly increasing owing to their ability to capture accurate and precise animal movements (He, Klarevas-Irby, Papageorgiou, Christensen, Strauss, and Farine 2022; Smith and Pinter-Wollman 2021; Cagnacci, Boitani, Powell, and Boyce 2010; Neethirajan and Kemp 2021). Since GPS devices are typically used to monitor a small proportion of individuals in a population, the network uncertainty resulting from such data is high (Farine and Strandburg-Peshkin 2015b). Therefore, missing data while performing social network analysis can have several implications, and the extent to which it is problematic depends on the nature and pattern of sampling strategies (Smith and Moody 2013; Smith and Morgan 2016; Smith, Morgan, and Moody 2022; Frantz, Cataldo, and Carley 2009). For animal social networks, performing statistical analysis without thorough information about the sources of bias and uncertainty could lead to incorrect inferences (Silk 2018; Gilbertson, White, and Craft 2021). If the animals are not sampled at random (which is often the case), the network metrics could be biased, potentially leading to inaccurate assessments of centrality, cohesion, or other network properties (Smith and Morgan 2016). The results may not accurately represent the true structure of the associations and impact the generalizability of the findings depending on the selection of nodes in the sample (Smith and Morgan 2016). Even if the sampling is done randomly, before performing social network analysis on a collection of GPS telemetry observations, it is essential to assess the appropriateness of the available data. It must be determined if the contacts resulting from such observations represent individual social preferences or are just chance encounters. The network measures obtained from a sample are the point estimates, reflecting the properties of the sample and not the whole population. The corresponding value that reflects the characteristics of the population might differ from this point estimate.

### 1.2. The five step workflow

(Kaur, Ciuti, Ossi, Cagnacci, Loison, Atmeh, McLoughlin, Reinking, Beck, Ortega, Kauffman, Boyce, and Salter-Townshend 2023) proposed a five-step workflow to assess the bias and uncertainty in the global and node-level metrics of animal social networks. The first step in the workflow is to determine whether the observations acquired by monitoring a sample of animals accurately reflect the true associations, screening out random encounters. For this, pre-network datastreams are permuted to form null networks, and the observed values of the network metrics of interest are compared to a null value distribution. In the second step, the sample’s robustness is evaluated by subsampling from the observed network, estimating uncertainty, and analysing bias in the retained network summary statistics. The third step estimates uncertainty in global network measurements by establishing confidence intervals around observed values using bootstrapping techniques. Node-level network metrics are affected by the proportion of individuals in the sample, therefore, the fourth step allows to assess the robustness of node-level network metrics using correlation and regression analyses. The final step generates confidence intervals for each node’s network metric value, enabling researchers to combine social connectedness with other ecological factors of interest (e.g., survival, mating strategy and success, habitat selection).

The R package aniSNA (Kaur 2024) is built around this five-step workflow and provides a platform to implement the statistical methods suggested as part of the workflow to assess the sufficiency of the data. Through the package aniSNA, we provide user-friendly functions to examine the suitability of a dataset concerning the research question at hand. The package functions are designed to work with GPS telemetry observations, however, the network structures obtained from other data sources can also be assessed for step two onwards of the five-step protocol presented by Kaur *et al*. (2023). The package’s functions allow users to apply bootstrapping techniques to obtain confidence intervals, which provide a measure of uncertainty around global and node-level network metrics. This is especially useful when a proportion of individuals are monitored from the population or the sampling proportion is unknown. In this paper, we present an overview of the package including the list of main functions. We then illustrate the step-wise workflow through an example of GPS telemetry observations of a large ungulate, the pronghorn (*Antilocapra americana*, Figure 1) and also discuss some of the applications of this workflow.

**Figure 1.**
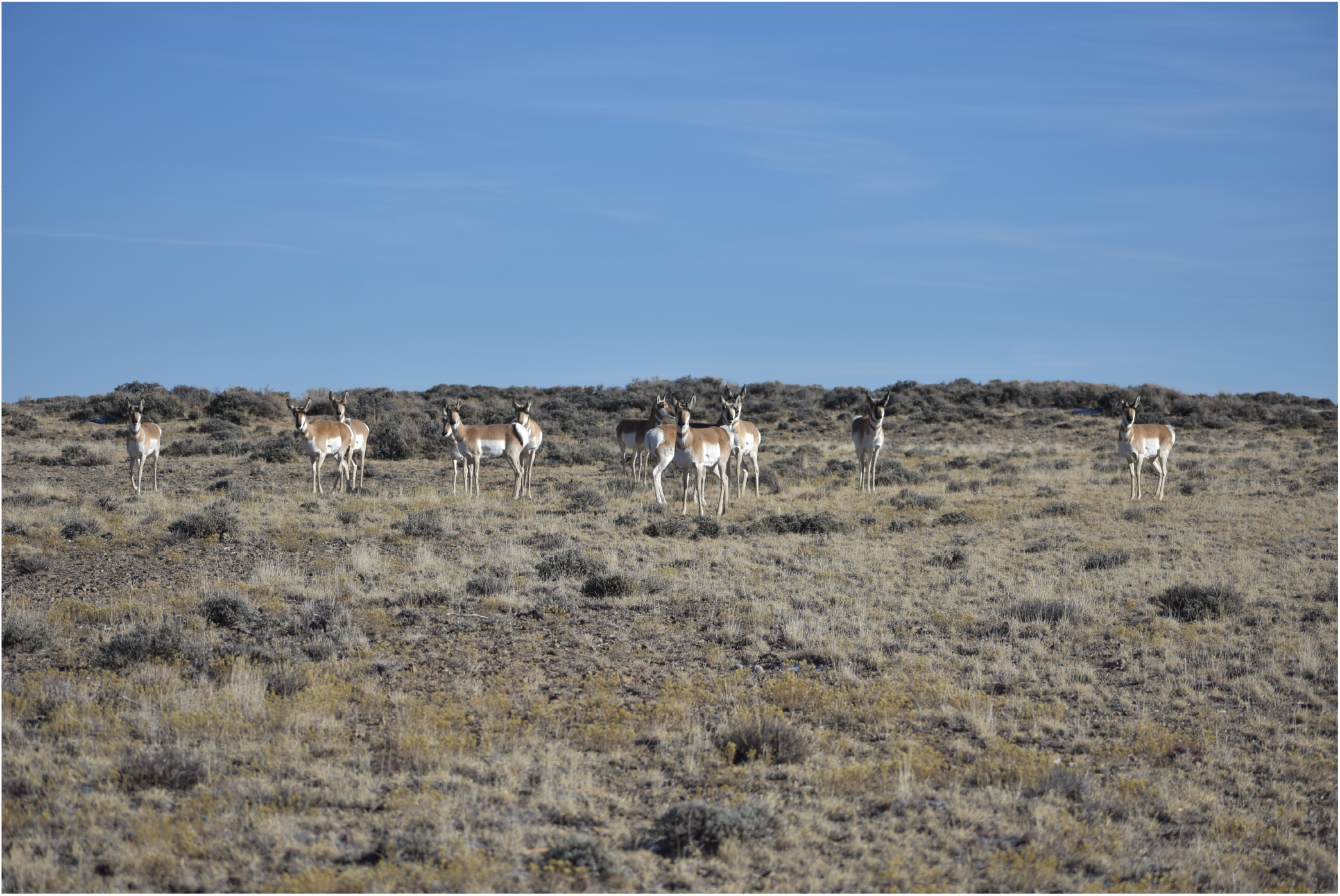
A group of female pronghorn from the study area (Image by Jacob D. Hennig).

## 2. aniSNA : Overview

aniSNA takes in a set of GPS telemetry observations of individuals from a population that has been monitored over a period of time. The data must consist of an individual animal ID, date and time of observation as well as the longitude and latitude coordinates of the location of observations. The input dataset needs to be an `R` dataframe consisting of four columns, namely “animal_id”, “datetime”, “latitude” and “longitude”. The latitude and longitude columns need to be in radians. A function called `get_coordinates_in_radian()` is available in the package, that can be used to convert degree coordinates into radians. This is a necessary step as radian values are later required to compute the distance between the individuals in order to consider interactions.

The data collected through GPS telemetry observations is used to obtain a network structure consisting of nodes and edges. The individuals that are monitored form the nodes of the network and an interaction between a pair of individuals form the edges (Farine and Whitehead 2015). A pair of individuals is deemed “associates” if they are observed within a predetermined temporal and spatial threshold. For instance, if two individuals are observed associating within a 25-meter radius within a 5-minute interval, as determined by the user, they are linked by an edge in the network. These network edges are assigned a weight, which is determined by the proportion of time the pair spends associating (He et al. 2022; Farine and Whitehead 2015; Kaur et al. 2023).

The networks generated through this method may either accurately depict the entire population or differ significantly from the true dynamics of the population, depending on various factors (James, Croft, and Krause 2009; Davis et al. 2018). These factors include the size of the population, the number and proportion of individuals monitored, the sampling strategy used to tag the individuals, the duration and frequency of observations, the ecological characteristics of the species, geographical features of the study area, among others (Davis et al. 2018; Kaburu, Balasubramaniam, Marty, Beisner, Fuji, Bliss-Moreau, and McCowan 2023; Gilbertson et al. 2021; Frantz et al. 2009; Silk 2018). Consequently, it’s essential to evaluate the quality of the data collected and the extent to which it accurately represents the population’s characteristics.

We now discuss the functions of the package aniSNA and provide a brief overview of each function’s purpose, usability, and outcomes. The functions in the package are organised around a workflow described as a five-step workflow by Kaur *et al*. (2023). A description of the main functions in aniSNA is provided in Table 1.

**Table 1:** A description of the main functions in aniSNA package.

| Function | Description |
| --- | --- |
| `bootstrapped_difference_pvalues` | Obtains two non-overlapping bootstrapped versions and generates p-values for the significance of difference. |
| `correlation_analyze` | Performs correlation analysis for node-level network metrics. |
| `get_coordinates_in_radian` | Converts latitude and longitude values from degree to radian. |
| `get_interactions` | Obtains interactions from raw GPS observations. |
| `get_network_summary` | Calculates and prints network summary statistics. |
| `get_spatial_threshold` | Calculates optimum spatial threshold for obtaining interactions from raw GPS observations. The threshold is obtained as the distance interval that captures maximum number of inter-individual interactions. |
| `global_width_CI` | Obtains width of confidence intervals using bootstrapped versions at each level of sub-sampling. |
| `global_CI` | Obtains confidence intervals for global network metrics. |
| `network_from_interactions` | Generates a network structure from interactions dataframe. |
| `node_level_CI` | Obtains confidence intervals for node-level network metrics. |
| `obtain_bootstrapped_samples` | Generates bootstrapped versions of a network's adjacency matrix. |
| `obtain_network_subsamples` | Generates subsamples of the observed network at a given level. |
| `obtain_permuted_network_versions` | Obtains permuted networks from raw datastream. |
| `plot_network` | Visualize animal social network. |
| `regression_slope_analyze` | Performs regression analysis for node-level network metrics. |
| `subsampled_network_metrics` | Generates subsamples of a network at a given level and obtains global network metrics of those subsamples. |
| `subsampled_permuted_network_metrics` | Generates subsamples of the permuted networks and obtain network metrics of those subsamples. |

## 3. Illustration

We illustrate the workflow of the functions in aniSNA with the help of a dataset consisting of GPS telemtry observations of Pronghorn (*Antilocapra americana*) `pronghorn_GPS_observations` (Reinking, Smith, Mong, Read, and Beck 2019). This large dataset consists of observations from a proportion of individuals sampled from the population of unknown total size and contains a unique animal identity number, date, time, and spatial coordinates of the observations. This collection of GPS telemetry observations is an example of a typical dataset that is available to the ecologists for analysis. Some of the important information such as the population size, exact sampling protocols are often unknown factors that affect the inference obtained from social network analysis (Sunga, Webber, and Broders 2021; Franks, Weiss, Silk, Perryman, and Croft 2020; Smith, Swain, Innocent, Nevison, and Hutchings 2019; Farine 2017). Through this dataset, we demonstrate how statistical inference can be obtained on the structure of the social network through the five-step workflow. We first load in the package and the dataset `pronghorn_GPS_observations in R`.

~~~
*R> library(aniSNA)
R> pronghorn_data <-read.csv(“pronghorn_GPS_observations.csv”)*
~~~

The dataset consists of GPS telemetry observations of 159 pronghorn, observed between November, 2013 and October, 2016 with each animal relocated every 2 hours. It is a dataframe consisting of four columns including a character column `animal_id`, a datetime column of class `POSIXct` representing calendar dates and times and latitude and longitude values in degrees. We obtain the radian values of longitude and latitude columns using the function `get_coordinates_in_radian()`.

~~~
*R> pronghorn_data <get_coordinates_in_radian(pronghorn_data)*
~~~

The above line of code introduces two additional columns in the existing dataframe namely `latitude_rad and longitude_rad`. We call the summary function on the dataset to ensure that all variables are in the expected format. It should be noted that the datetime column should be in `POSIXct` class, if not, it can be converted to it using the function `as.POSIXct()`.

~~~
*R> pronghorn_data$datetime <-as.POSIXct(pronghorn_data$datetime*,
                                           *format = “%Y-%m-%d %H:%M:%OS”)
R> summary(pronghorn_data)*
~~~

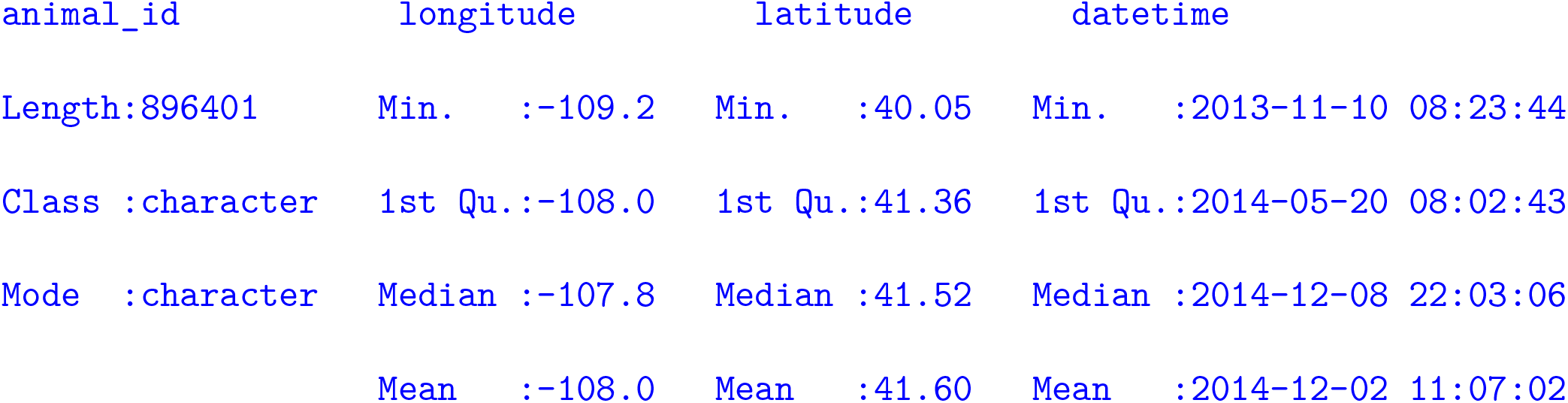

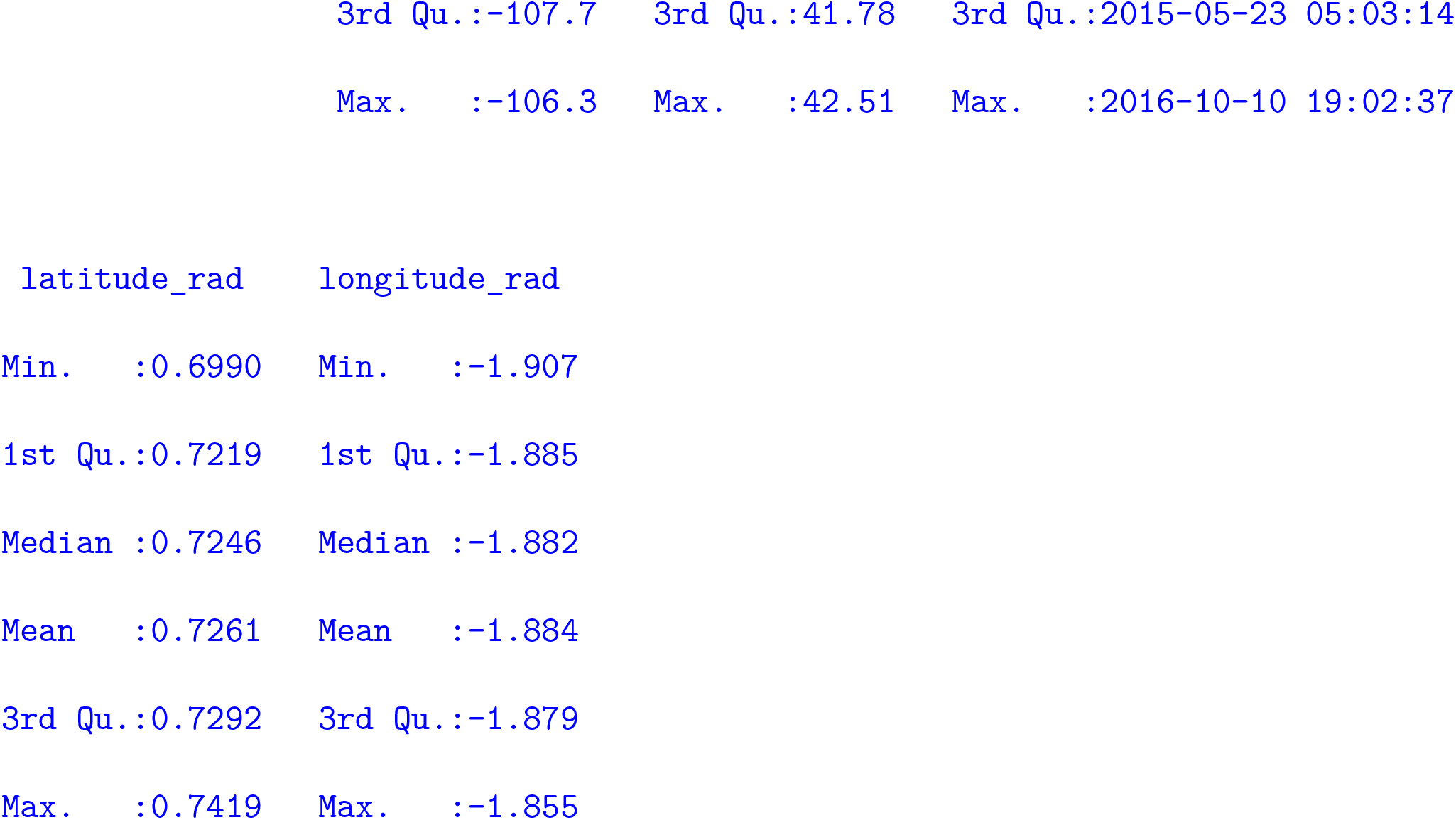

### Identify interactions and form network structure

Now we use this dataset to identify interactions. We choose a spatial and temporal threshold to obtain pairs of interacting individuals. The spatial threshold defines the maximum distance in metres within which two animals are considered interacting (Davis et al. 2018). This value can be dictated by the prior information on species ecology and the research question. We can also identify an optimum value of spatial threshold from the raw set of GPS observations. The optimum value will be the shortest distance threshold that captures maximum number of true interactions within all pairs. This method is also called identifying the first mode of number of interactions and is as per the suggestions by He *et al*. (2022) and Kaur *et al*. (2023).

To do that, we must first determine the greatest distance within which we may reasonably expect to discover the ideal spatial threshold. For pronghorn, we specify this distance to be 50m and retrieve all possible interactions within this spatial threshold by using the function `get_interactions()` from the package. Next, we determine the ideal spatial threshold value using the function `get_spatial_threshold()`. The function takes an argument `interval_size` which represents the size of the bins within which we are searching the optimal value. Note that this function allows for parallel processing and the user can specify the number of cores if multiple cores are available.

~~~
*R> pronghorn_50m_interactions <-get_interactions(pronghorn_data*,
                              *temporal_thresh = 7*,
                              *spatial_thresh = 50*,
                              *n_cores = 3)
R> get_spatial_threshold(pronghorn_50m_interactions, interval_size = 1)*
13
~~~

The function `get_spatial_threshold()` yields a result of 13 when the interval size is one, indicating that 13 metres is the ideal value to use for spatial threshold. In other words, our routine suggests that in this specific case, based on the observed data, the two animals are considered to be interacting, when they are within 13 meters of each other. We selected a temporal threshold of 7 minutes in order to get interactions using the function `get_interactions()`. The time interval when two animals seen within a specific distance are deemed to be interacting is known as the temporal threshold (See Kaur *et al*. (2023) for more details). Basically, this allows for some animals that are together but are relocated at slighlty different times (e.g. 11:57 am and 12:01 pm - quite typical in GPS telemetry) to be deemed together (in our example we picked 7 minutes but users can change it). With the ideal spatial and temporal thresholds, we can now acquire the complete set of interactions. Again, the package function `get_interactions()` is used.

~~~
*R> pronghorn_interactions <-get_interactions(pronghorn_data*,
                                 *temporal_thresh = 7*,
                                 *spatial_thresh = 13*,
                                 *n_cores = 3)*
~~~

This produces a dataframe consisting of five columns and 8254 observations. The first two columns, “Animal_A” and “Animal_B” respectively, contain the animal IDs of two animals in an interacting pair, relocated together within 13 meters and 7 minutes. Columns three and four contain the observation timestamps for each of the animals in the pair and last column contains the observed distance in metres between that pair. The Euclidean distance between the geographic coordinates of two individuals at that particular time is used to compute the distance.

In the next step, we obtain a network structure from this set of interactions. This is accomplished by using the package function `network_from_interactions()`, which yields an igraph object. The function `plot_network()` may be used to display the network structure. A visualisation of the pronghorn network obtained by this function is given in Figure 2. In this figure, two dense groups are easily discernible. Please take note that this representation only offers a rudimentary understanding of the structure and is barely informative.

**Figure 2.**
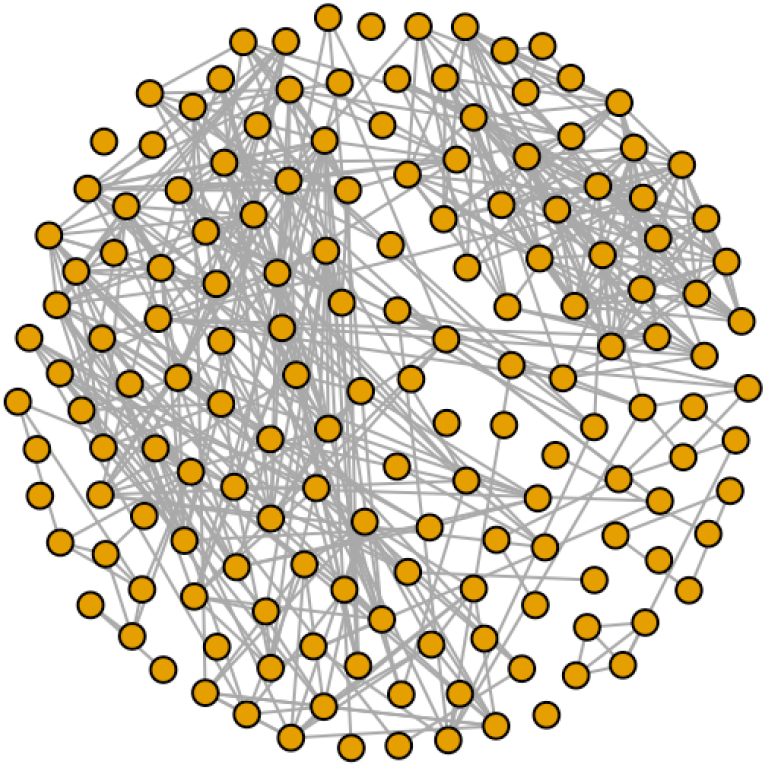
Network visualisation obtained using plot_network() function.

~~~
*R> pronghorn_network <-network_from_interactions(pronghorn_data*,
                                             *pronghorn_interactions*,
                                             *n_cores = 3)
R> plot_network(pronghorn_network)*
~~~

We use the function `get_network_summary()` to investigate network properties and generate network metric values for the pronghorn network obtained from the available sample.

~~~
R> get_network_summary(pronghorn_network)
~~~

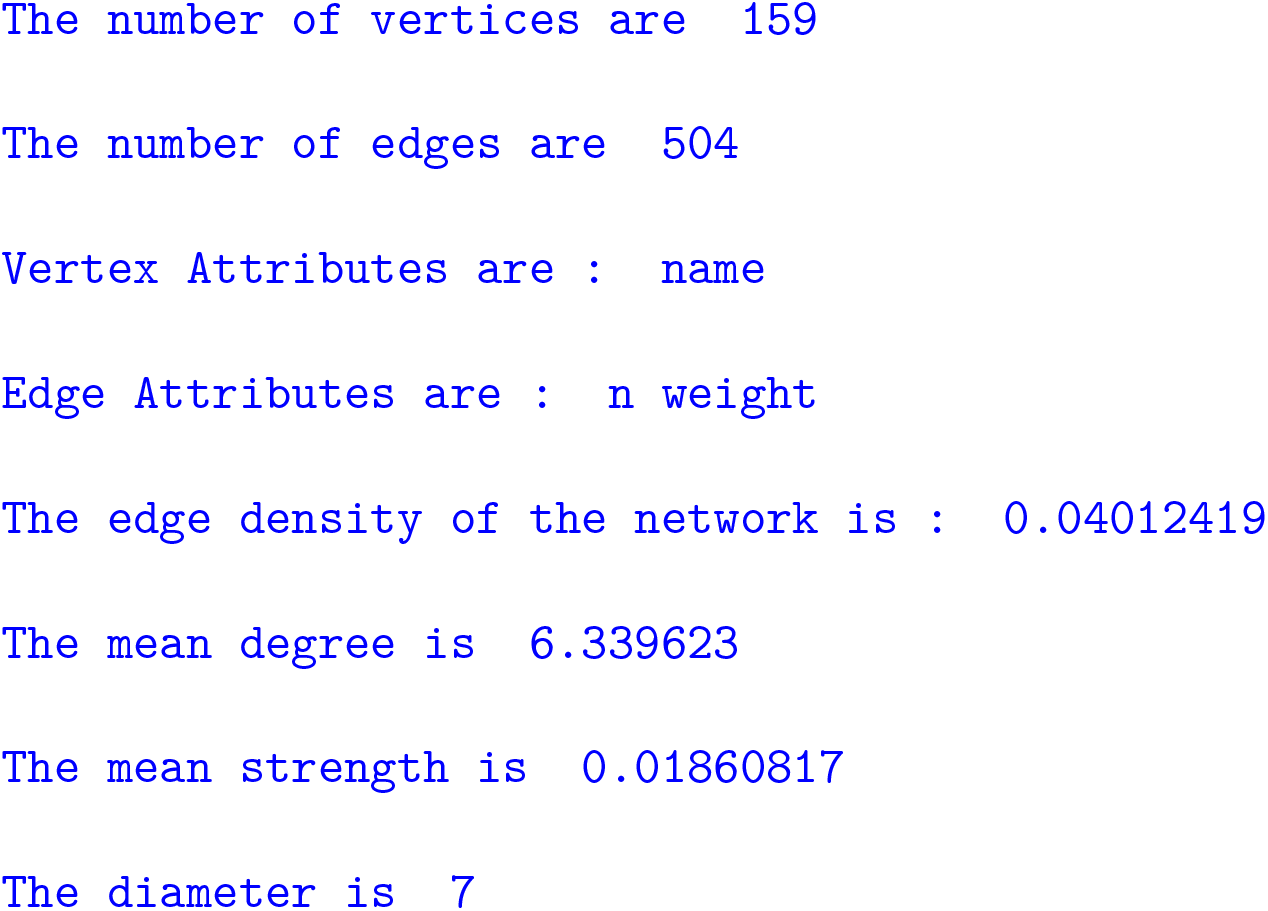

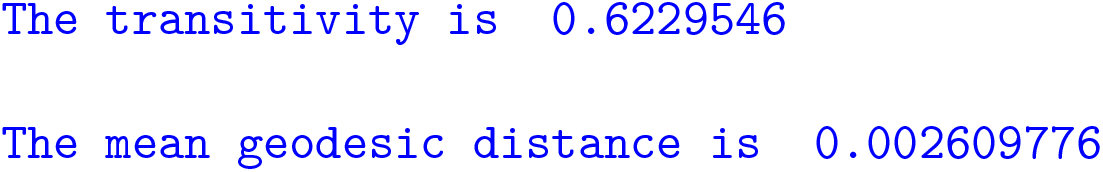

At this stage, the network structure is ready for the analysis using the five-step workflow (Kaur et al. 2023). A summary of the steps so far is provided in Figure 3.

**Figure 3.**
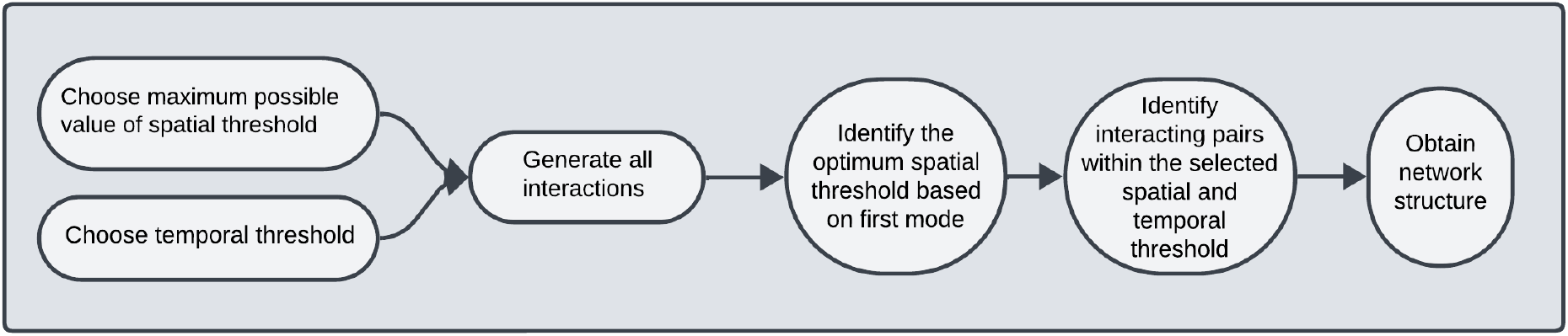
A flowchart of steps for network formation.

#### Step 1 : Pre-network data permutations

Social network analysis for animal communities is increasingly widely used (Silk et al. 2017; Hock and Fefferman 2011; Webber, Schneider, and Wal 2020; Pinter-Wollman, Hobson, Smith, Edelman, Shizuka, De Silva, Waters, Prager, Sasaki, Wittemyer, Fewell, and McDonald 2013). Typically, researchers focus on specific study questions and seek evidence to test theories. Consequently, they choose the most suitable network metric to represent a network feature and aid in making inferences (Sosa, Sueur, and Puga-Gonzalez 2021). Therefore, the first step is to determine if the observed interactions and the resulting network metric captured by the observed sample are indeed caused by social preferences or if those are the results of random encounters (Croft, Madden, Franks, and James 2011; Sundaresan, Fischhoff, and Dushoff 2009; Farine 2014). In this reference, null models are created to account for the non-social factors that lead to animal co-occurrence (Farine 2017; Spiegel, Leu, Sih, and Bull 2016).

High-resolution GPS telemetry data often generates autocorrelated streams, with the extent of autocorrelation varying based on individual speed. To maintain the autocorrelation structure of individual movements but randomize contacts, pre-network datastream permutations are obtained. This methodology segments daily tracks for each individual and shuffled their dates, ensuring unaffected home ranges of animals in the permuted data but randomizing contacts in the null model (Farine and Carter 2022; Farine 2017; Spiegel et al. 2016).

The underlying purpose is to determine if the available data is appropriate to answer the research question in consideration. With this aim, we compare the observed network metric to a distribution of network metrics derived from permuted data (See Kaur *et al*. (2023) for more details). Permuted data represents observations that have no underlying structure and which would have been observed if the animals moved arbitrarily without any social or other preferences (Farine 2017).

As a result, it is critical to determine if the given data accurately captures the aspect of the population that is being examined. We use seven global network metrics commonly employed in animal social network studies including edge density, mean strength, diameter, transitivity, assortativity degree, modularity, and global efficiency (Kaburu et al. 2023; Shimada and Sueur 2014). Each of these network metrics represents a particular aspect of the animal social network (Sosa et al. 2021) and are summarised in Table 2.

**Table 2:** Global network metrics used in the analysis of pronghorn network.

| Global Network Metric | What does the metric measure? |
| --- | --- |
| Edge Density | The proportion of completed edges in the network. |
| Transitivity | The amount of clustering in the network, calculated as a function of completed triangles relative to possible triangles. |
| Diameter | The shortest distance between the two most distant nodes in the network. |
| Modularity | This measure quantifies the degree of segregation or partitioning in the network structure. |
| Mean Strength | The average strength of a node in the network. |
| Assortativity Degree | This metric measures the tendency of nodes in the network to connect to other nodes with similar degree values. A positive assortativity degree indicates that high-degree nodes are more likely to connect to other high-degree nodes, while low-degree nodes tend to connect to other low-degree nodes. |
| Global Efficiency | This measure quantifies the efficiency of information transfer within the entire network and is calculated as the average inverse shortest path length between all pairs of nodes in the network. |

The package function `obtain_permuted_network_versions()` is used to generate permuted versions. The first argument to the function is the dataframe of raw GPS telemetry observations, and the second and third arguments are the temporal and spatial thresholds. The user also needs to specify the number of permuted versions that should be generated. The function has a default value of 100 permutations and we can specify the number of cores to allow for parallel processing. Note that this function is computationally intensive and the users are suggested to run this on a remote server if possible. The time taken depends on the number of observations in the raw GPS data set and the number of permutations required. To obtain 100 permutations for the dataframe `pronghorn_data` using four cores, it took approximately four hours.

~~~
*R> pronghorn_permutations <-obtain_permuted_network_versions(pronghorn_data,
                                          temporal_thresh = 7,
                                          spatial_thresh = 13,
                                          n_permutations = 100,
                                          n_cores = 4)
R> plot(pronghorn_permutations, pronghorn_network,
          network_metrics_functions_list =
                      c(“Edge density” = function(x) igraph::edge_density(x),
                      “Mean strength” = function(x) mean(igraph::strength(x)),
                      “Diameter” = function(x) igraph::diameter(x, weights = NA),
                      “Transitivity” = function(x) igraph::transitivity(x),
                      “Assortativity degree” = function(x)
                           igraph::assortativity.degree(x),
             “Modularity” = function(x) igraph::modularity(x,
                     igraph::membership(igraph::cluster_walktrap(x)),
                           weights = igraph::E(x)$weight),
             “Global efficiency” = function(x) igraph::global_efficiency(x)))*
~~~

The function `obtain_permuted_network_versions()` returns a list of size `n_permutations` in which each element is an `igraph` network obtained from the permuted versions of the raw data stream. The returned list belongs to class `list_permuted_networks`. The function `plot()` is used on the returned list of networks which obtains a visualisation of network metrics histogram (see Fig 4). The user can specify the network metrics that are of interest in the form of a list. In this example, we have picked seven network metrics and defined their functions. This feature enables the user to assess any network measure of interest by giving a simple function definition for it. The plots obtained in this way also indicate the value of the network metric in the observed network. The user can compare the position of the observed network metric with respect to the samples of network metrics obtained from permuted versions of raw data. Using this visualisation, it can be verified if the data captures non-random aspects of the network. Note that the user is free to add new network metrics, delete them from this list, or use a different set of network metrics that is better suited to the research question.

**Figure 4.**
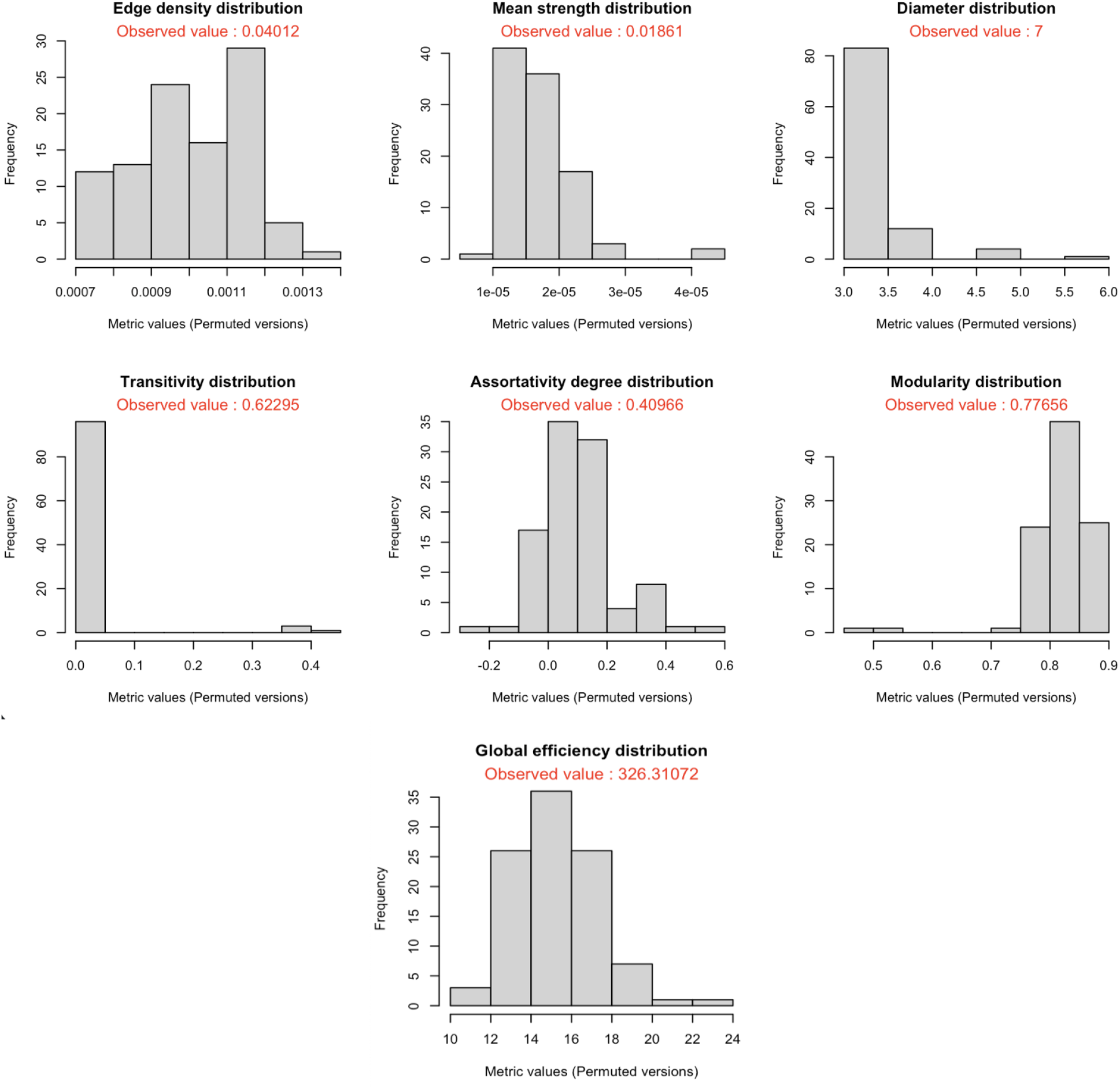
Histogram of the network metric values obtained from permuted versions of the raw data stream. If the observed value falls within the 95% confidence interval, the network metric should not be used since it does not account for that non-random aspect of the associations.

~~~
*R> assortativity_observed <-igraph::assortativity.degree(pronghorn_network)
R> print(assortativity_observed)*
          0.4096592
*R> assortativity_degree_null_values <-unlist(lapply(pronghorn_permutations*,
                         *function(x) igraph::assortativity.degree(x)))
R> round(quantile(assortativity_degree_null_values*,
          *probs = c(0.025, 0.975)), 5)*
  2.5% 97.5%
-0.09091 0.38944
*R> modularity_observed <-igraph::modularity(pronghorn_network*,
                    *igraph::membership(
                    igraph::cluster_walktrap(pronghorn_network))*,
                    *weights = igraph::E(pronghorn_network)$weight)
R> print(modularity_observed)*
  0.7765574
*R> modularity_degree_null_values <-unlist(lapply(pronghorn_permutations*,
          *function(x)
          igraph::modularity(x*,
          *igraph::membership(
          igraph::cluster_walktrap(x))*,
          *weights = igraph::E(x)$weight)))
R> round(quantile(modularity_degree_null_values*,
                         *probs = c(0.025, 0.975)), 5)*
   2.5% 97.5%
0.74521 0.88853
~~~

From the visualisations obtained in this case, we see that the observed values of assortativity degree and modularity lie within the distribution of null values. We further investigate if the observed values for these two network metrics lie within the middle 95% in the distribution of null values. For this, we use the `quantile()` function to extract 95% confidence intervals. For assortativity degree, the observed value does not lie within 95% confidence interval of null values but for modularity, it does. As the network metric modularity is used to assess the presence of community structure in networks, this indicates that the sample that we have has a pattern of clustering among nodes which could have been present in any random network of associations. It does not reveal any underlying patterns of organization in the pronghorn population based on the available sample. Therefore, modularity should not be used for further analysis such as hypothesis testing on the pronghorn dataset. In general, if the observed value falls outside the 95% CI then the data collected does capture a social structure that differs from random associations. Therefore we accept the network metric assortativity degree along with the rest of the five network metrics for further analysis.

#### Step 2a : Subsampling from the observed sample

In the second phase of the workflow, the goal is to determine if the network metrics chosen at this stage remain stable and robust under sampling and understand the level of bias as the sample size lowers. To do so, we perform subsampling and assess change in the network metrics’ values with decreasing sample size. This provides an idea of the extent of bias that can be expected in the values of these network metrics as the observed datasets are the subsets of the population. The package function `subsampled_network_metrics()`enables users to construct several subsamples at different proportions. The function selects the nodes at random without replacement and generates a network from the selected nodes. All interactions within the specified nodes are retained, while the remainder are discarded. This procedure mimics the process of random sampling from the population and depicts the network structure if initially an even lower proportion of the population was randomly sampled. In this example, we proceed with the six network metrics chosen in step 1 and supply those as a list to the function argument `network_metrics_functions_list`.

~~~
*R> pronghorn_subsampling <-subsampled_network_metrics(pronghorn_network*,
         *n_simulations = 100*,
         *subsampling_proportion = c(0.1, 0.3, 0.5, 0.7, 0.9)*,
         *network_metrics_functions_list =
  c(“Edge density” = function(x) igraph::edge_density(x)*,
  *“Mean strength” = function(x) mean(igraph::strength(x))*,
  *“Diameter” = function(x) igraph::diameter(x, weights = NA)*,
  *“Transitivity” = function(x) igraph::transitivity(x)*,
  *“Assortativity degree” = function(x)
  igraph::assortativity.degree(x)*,
  *“Global efficiency” = function(x) igraph::global_efficiency(x)
  )
)
R> plot(pronghorn_subsampling, pronghorn_network*,
                 *network_metrics_functions_list =
    c(“Edge density” = function(x) igraph::edge_density(x)*,
    *“Mean strength” = function(x) mean(igraph::strength(x))*,
    *“Diameter” = function(x) igraph::diameter(x, weights = NA)*,
    *“Transitivity” = function(x) igraph::transitivity(x)*,
    *“Assortativity degree” = function(x)
             igraph::assortativity.degree(x),
    “Global efficiency” = function(x) igraph::global_efficiency(x)
     )
)*
~~~

The function `subsampled_network_metrics()` returns a list of length equivalent to the number of metrics passed in the argument `network_metrics_functions_list`. The list belongs to the class “Subsampled_Network_Metrics” which allows the user to use the plot function to obtain visualisations corresponding to each network metric. The visualisation consists of boxplots (Figure 5) that represent the distribution of network metric values obtained from multiple subsamples at each level of subsampling. Edge density, transitivity, and assortativity degree remain unbiased even when the subsampling proportion is lowered to 30%. On the other hand, the network metrics mean strength, diameter, and global efficiency display a biased behaviour as the number of nodes in the subsample decreases. Note that the diameter tends to plateau at 8 depicting that even if we had a larger sample from the population, the diameter would remain 8 or very close to it. Depending on the study objective, the user must decide if the chosen network metrics are compatible with their aims based on their potential to exhibit biased behaviour. Inferences based on the network metrics that tend to get biased as the sample size lowers should be made carefully. When the sampling proportion is unknown, conclusions on the population should not be made based on the sample for the metrics displaying biased behaviour.

**Figure 5.**
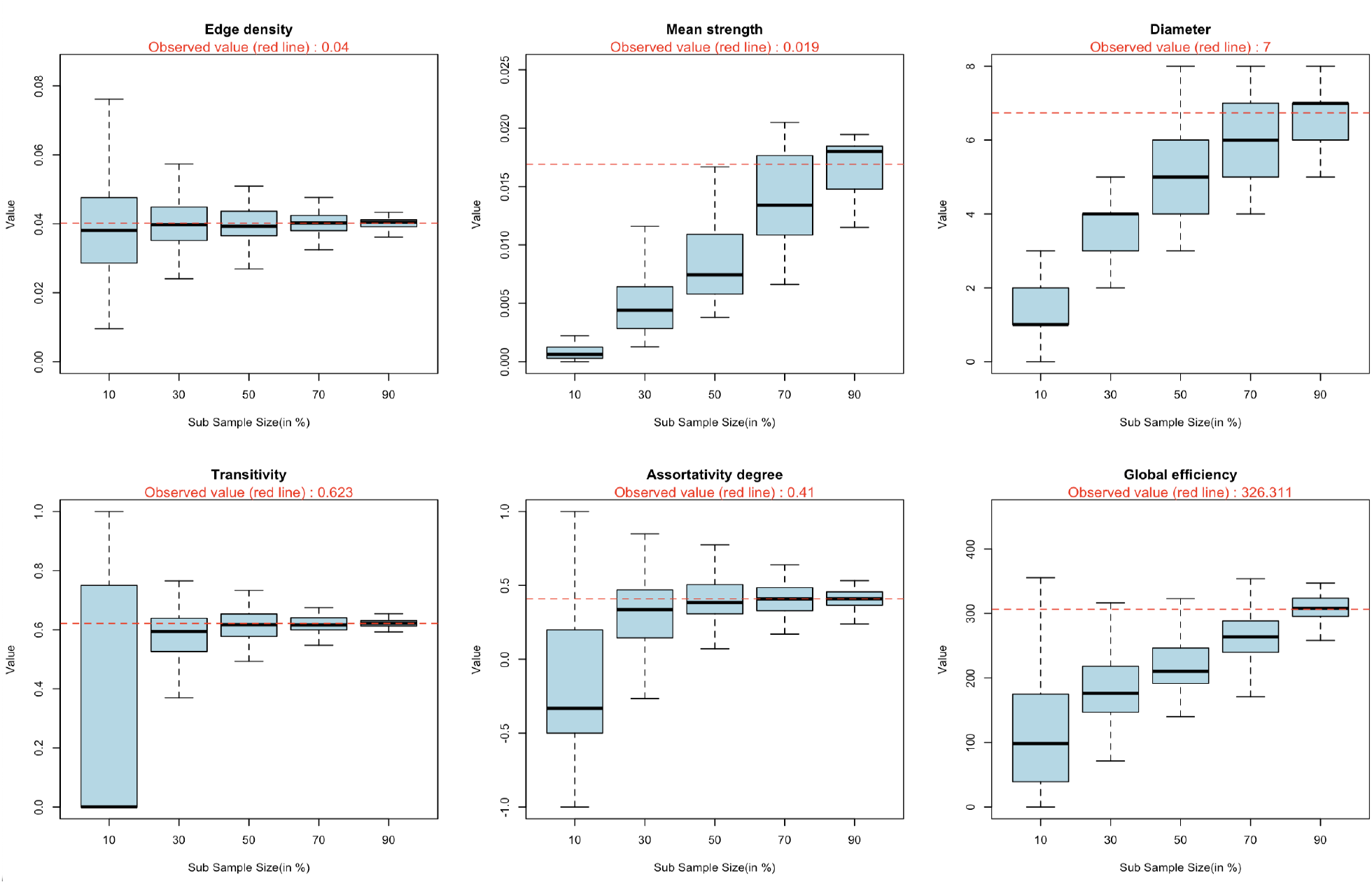
Effect of sub-sampling on six global network metrics. The horizontal red line in each plot represents the metric value in the observed network. The boxplots denote the distribution of network metric values obtained from the observed networks by taking 100 sub-samples at each level.

#### Step 2b : Comparing subsamples of the observed and the permuted networks

The package also allows the user to compare the subsamples from the observed network to those obtained from permuted versions of the observed network. This comparison allows to understand under what level of sampling would the current non-random metrics resemble the random values (Kaur et al. 2023). The function `subsampled_permuted_network_metrics()` is used to obtain the values at each level of subsampling and returns an object of class “Subsampled_Permuted_Network_Metrics”. The object can then be passed to plot() to obtain a visualisation that implements the plots shown in Figure 6.

**Figure 6.**
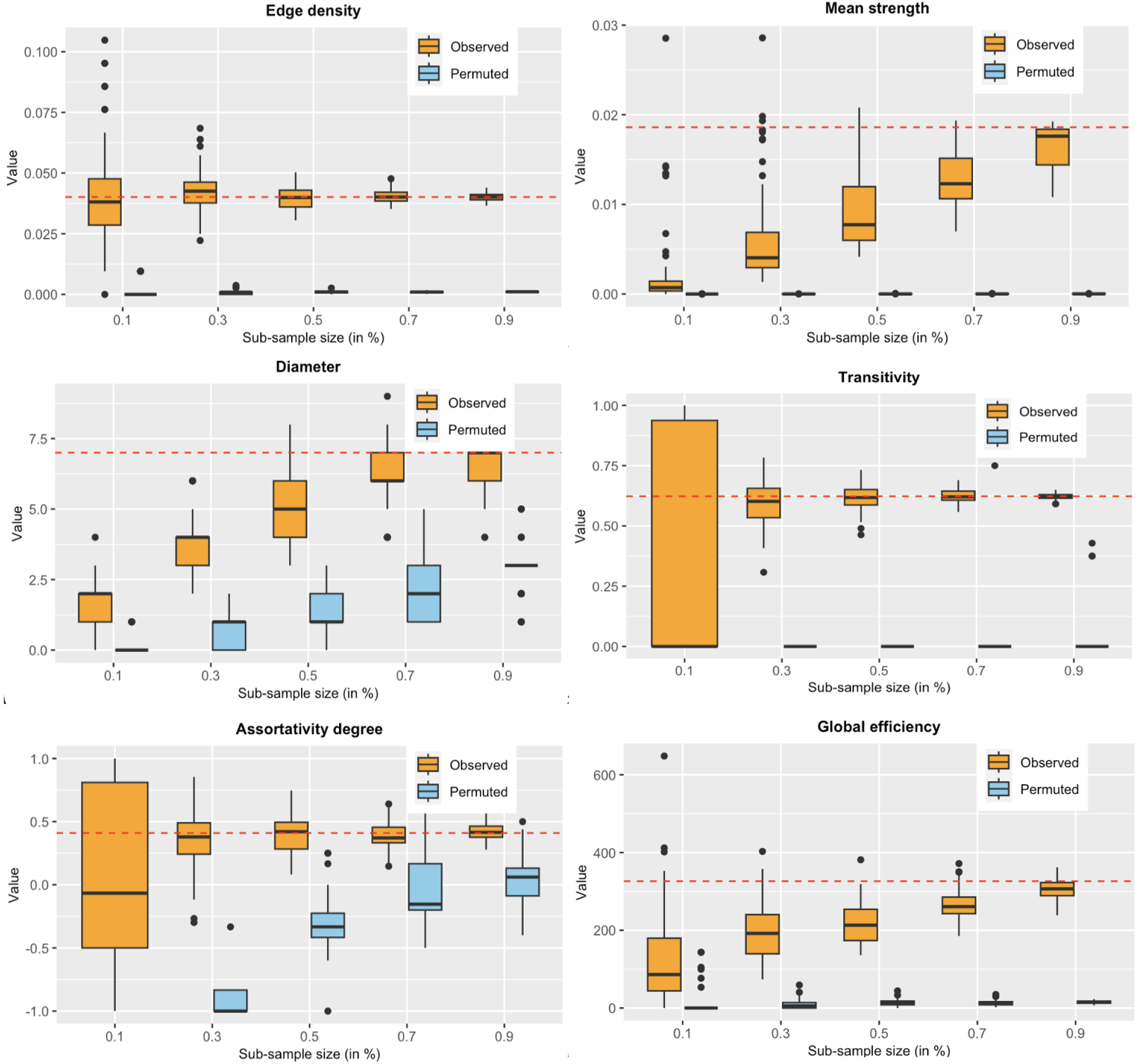
Comparison of the subsamples of the observed network to those of permuted net-works. The blue boxplots are obtained by calculating network metric values for 1000 permuted versions. The orange boxplots are the ones that we obtained from subsampling the observed network. The horizontal red line in each plot represents the observed metric value. Comparing the subsamples of the observed network to those of permuted networks identifies the sampling proportion where the non-random aspects of the observed network resemble to those of random networks.

~~~
*R> permuted_subsamples_comparison <-subsampled_permuted_network_metrics(
      pronghorn_permutations,
      subsampling_proportion = c(0.1, 0.30, 0.50, 0.70, 0.90),
      network_metrics_functions_list =
  c(“Edge density” = function(x) igraph::edge_density(x),
     “Mean strength” = function(x) mean(igraph::strength(x)),
     “Diameter” = function(x) igraph::diameter(x, weights = NA),
     “Transitivity” = function(x) igraph::transitivity(x),
     “Assortativity degree” = function(x) igraph::assortativity.degree(x),
     “Global efficiency” = function(x) igraph::global_efficiency(x))
)
R> plot(permuted_subsamples_comparison,
   pronghorn_network,
   network_metrics_functions_list =
   c(“Edge density” = function(x) igraph::edge_density(x),
   “Mean strength” = function(x) mean(igraph::strength(x)),
   “Diameter” = function(x) igraph::diameter(x, weights = NA),
   “Transitivity” = function(x) igraph::transitivity(x),
   “Assortativity degree” = function(x) igraph::assortativity.degree(x),
   “Global efficiency” = function(x) igraph::global_efficiency(x))
)*
~~~

For the network metrics edge density and transitivity, even if 10% of the nodes were sampled from the population, the observed network would still be very different to a network of random interactions. For the network metric assortativity degree, the overlapping between the two boxplots starts at 90% sampling level. However, for the permuted network versions, assortativity degree becomes highly biased with decrease in the sub-sample size. For the sample of pronghorn population, the network metrics edge density, mean strength, transitivity and global efficiency are captured very well and the network could be differentiated from a random network even at lower sample sizes. This result can help determine the minimum level of sampling required to obtain correct inferences for future research studies, given the research question requires analysis of these network metrics.

#### Step 3 : Estimate uncertainty around the point estimates of global network metrics

One way of estimating uncertainty around the network metrics’ point estimates is to obtain confidence intervals (Snijders and Borgatti 1999). Animal data is inherently variable and confidence intervals acknowledge and account for this variability, helping to avoid overconfidence in the precision of network metric estimates (Whitehead 2009; Borgatti, Everett, and Freeman 2014).

In animal social network studies, care should be taken while interpreting the confidence intervals obtained for the observed network metrics Farine and Carter (2022); Lusseau, Whitehead, and Gero (2008); Whitehead (2008). In step 2 of the workflow, the subsampling analysis identified the network metrics that remain unbiased as the sample size lowers. For such network metrics, confidence intervals provide a range of values within which the true population parameter is likely to fall. However, for the network metrics that became unbiased with lowering sample size such as mean strength, diameter, and global efficiency, confidence intervals obtained for the observed value may not contain the true population parameter. Therefore, inference using confidence intervals for the full population should not be made using a sample for such network metrics.

However, confidence intervals can provide other useful information even for biased network metrics. It can be used to test the significance of the difference between networks obtained from two samples of the same size. For example, if a researcher is interested in testing hypotheses such as the difference in mean strength of the sampled network in summer and winter. Confidence intervals allow for a direct comparison of the estimated network metrics. If the intervals for the two groups do not overlap, it suggests a statistically significant difference.

In general, confidence intervals provide a more comprehensive picture of the network structure compared to point estimates alone. Instead of relying solely on a single value, it conveys the range of plausible values for a network metric. This enables researchers and decision-makers to understand the level of uncertainty associated with their social network estimates which is particularly important when decisions are based on statistical inference. Kaur *et al*. (2023) described the importance and need to obtain confidence intervals around the point estimates of network metrics generated from an observed sample from a population and suggested to implement it as the third step in the five-step workflow. In aniSNA, the function `global_CI()` generates 95% confidence intervals around the observed network metric estimates as shown in the code below.

~~~
*R> pronghorn_global_CI <-global_CI(
          pronghorn_network,
          n_versions = 100,
         network_metrics_functions_list =
   c(“Edge density” = function(x) igraph::edge_density(x),
   “Mean strength” = function(x) mean(igraph::strength(x)),
   “Diameter” = function(x) igraph::diameter(x, weights = NA),
   “Transitivity” = function(x) igraph::transitivity(x),
   “Assortativity degree” = function(x) igraph::assortativity.degree(x),
   “Global efficiency” = function(x) igraph::global_efficiency(x))
 )
R> round(pronghorn_global_CI,4)*
~~~

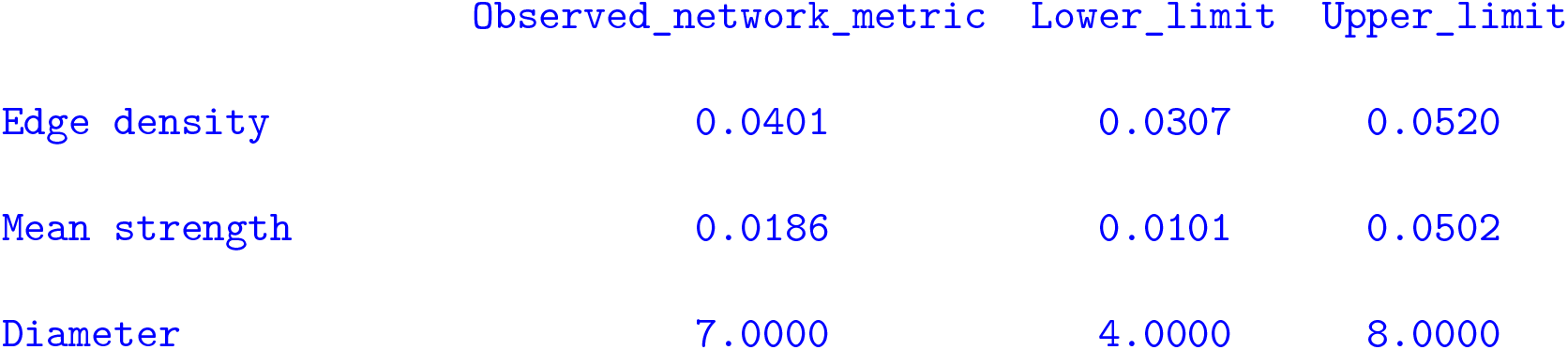

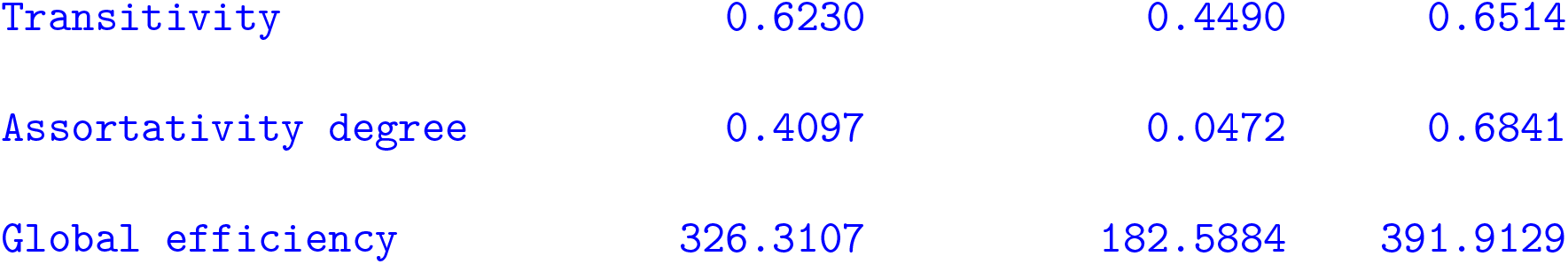

Confidence intervals also indicate the precision of the estimated network metric with respect to the size of the sample. A narrower interval implies a more precise estimate, while a wider interval suggests greater uncertainty. The package function `global_width_CI()` allows to investigate how the width of confidence intervals around the point estimates of global network metrics change as the sample size is lowered. The function `global_width_CI()` takes in the observed network as the first argument. The user can specify the number of bootstrapped versions that should be used to obtain confidence intervals (See Kaur *et al*. (2023) for more details on bootstrapping.) The argument `n.iter` represents the number of iterations at each level of sub-sampling over which the mean of confidence intervals is calculated.

~~~
*R> pronghorn_width_CI <-global_width_CI(pronghorn_network,
    network_metrics_functions_list =
    c(“Edge density” = function(x) igraph::edge_density(x),
    “Mean strength” = function(x) mean(igraph::strength(x)),
    “Diameter” = function(x) igraph::diameter(x, weights = NA),
    “Transitivity” = function(x) igraph::transitivity(x),
    “Assortativity degree” = function(x) igraph::assortativity.degree(x),
    “Global efficiency” = function(x) igraph::global_efficiency(x))
 )
R> plot(pronghorn_width_CI)*
~~~

The function `global_width_CI()` returns a list of vectors of class “Width_CI_matrix” which the user can pass into the `plot()` function and obtain a visualisation as shown in Figure 7. As the sample size is lowered, the uncertainty in the observed network metric increases, and therefore, the width of the confidence intervals widens. However, for some of the network metrics like mean strength and diameter, the width of confidence intervals declines when the sub-sample size is lowered. This is because for these network metrics, the number of nodes in the network sample has a direct impact on the observed values of these metrics (See Kaur *et al*. (2023) for more details) and therefore, a scaled version for these metrics should be considered. In the function `global_width_CI()`, the user has a choice to specify the network metrics whose scaled versions need to be considered. The network metrics that are directly affected by the number of nodes present in the sample should be specified here. We repeat this analysis by specifying these metrics.

**Figure 7.**
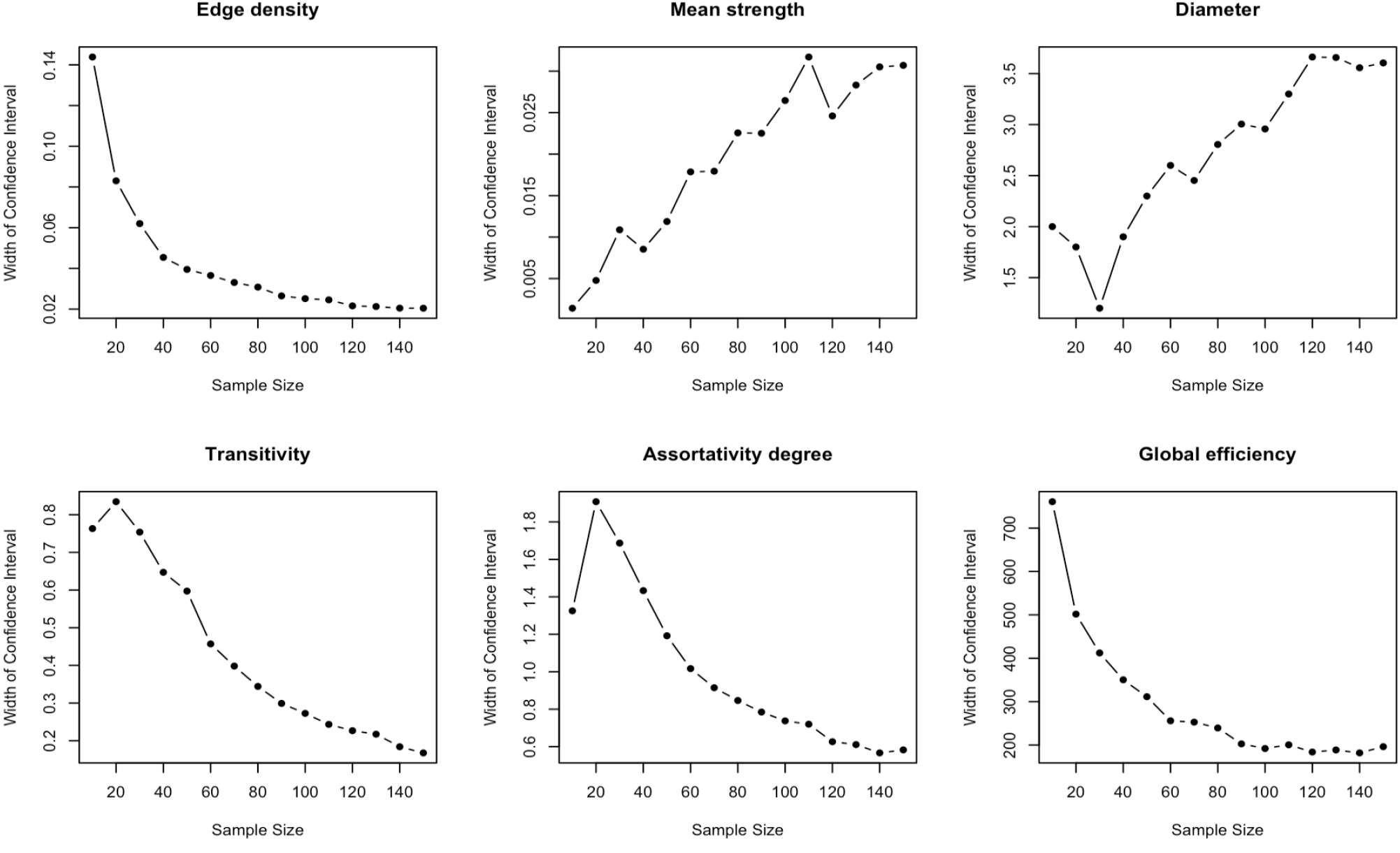
The plots show the mean widths of 95% confidence intervals obtained from boot-strapped sub-samples of a network. The x-axis indicates the number of nodes in the sample and y-axis denotes the mean width of confidence intervals. Except for mean strength and diameter, the mean widths of all network metrics increase with lower sample size indicating increasing uncertainty around the point estimate of the network metrics. As the values for mean strength and diameter are directly affected by the number of nodes present in the network, we consider scaled versions of these two metrics such that the values at each level are scaled by the number of nodes at that level.

~~~
*R> pronghorn_width_CI_scaled <-global_width_CI(pronghorn_network,
     network_metrics_functions_list =
     c(“Edge density” = function(x) igraph::edge_density(x),
     “Mean strength” = function(x) mean(igraph::strength(x)),
     “Diameter” = function(x) igraph::diameter(x, weights = NA),
     “Transitivity” = function(x) igraph::transitivity(x),
     “Assortativity degree” = function(x) igraph::assortativity.degree(x),
     “Global efficiency” = function(x) igraph::global_efficiency(x)),
     scaled_metrics = c(“Mean strength”, “Diameter”)
  )
R> plot(pronghorn_width_CI_scaled)*
~~~

We investigate the scaled versions of these network metrics by obtaining the corresponding plots for these (Figure 8). After scaling, the plot for the network metric diameter shows similar behaviour to that of unscaled versions of other network metrics. Interestingly, the confidence interval widths of mean strength first increase and then decrease as the sample size is lowered. This indicates that when the sample size falls below a specific threshold, the observed network splits into smaller disjoint sub-networks, and even in bootstrapped versions of these smaller networks, the mean strength values remain low.

**Figure 8.**
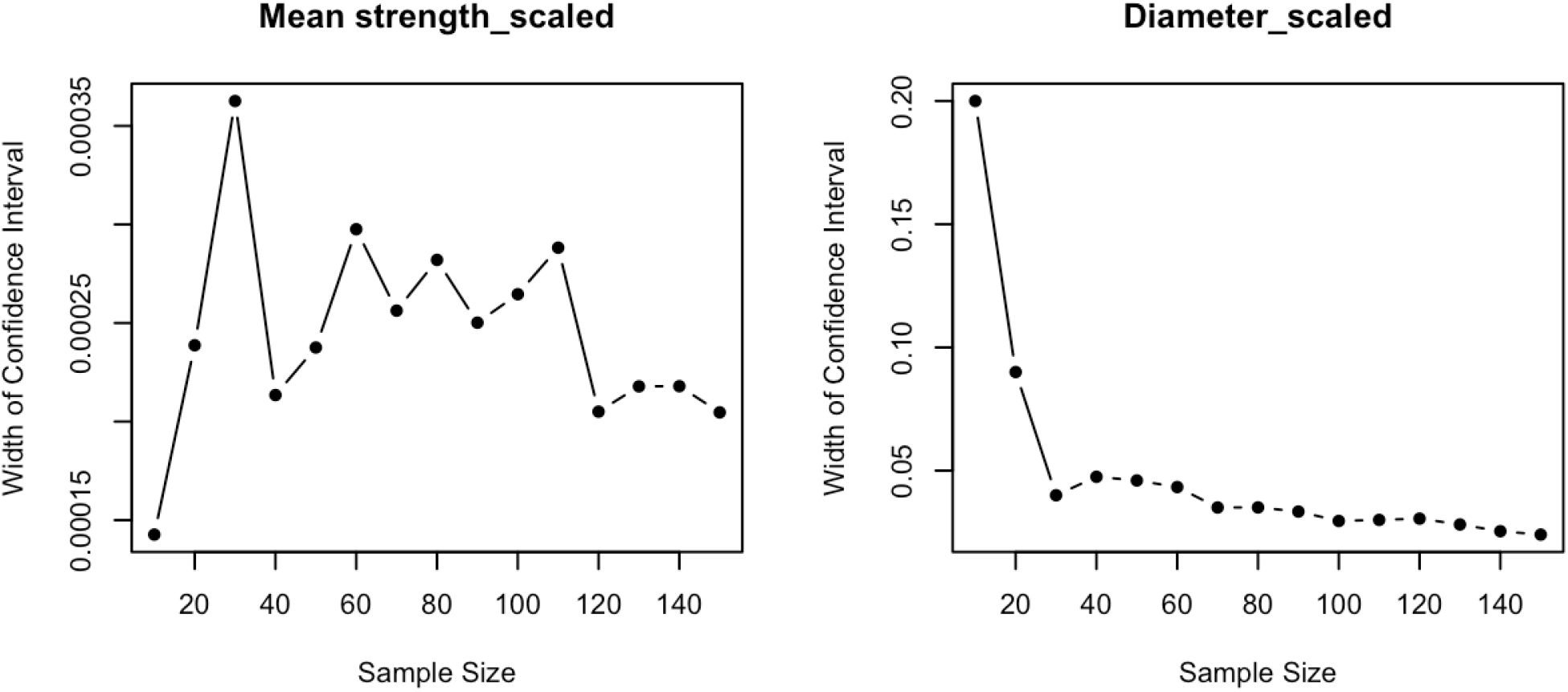
The plots show the mean widths of 95% confidence intervals for the scaled versions of the chosen network metrics. The x-axis indicates the number of nodes in the sample and y-axis denotes the mean width of confidence intervals.

#### Step 4a : Correlation analysis between node-level network metrics of observed and smaller subsamples

The fourth step in the five-step workflow (Kaur et al. 2023) concerns the node-level network metrics. We use eight node-level network metrics commonly used in animal social network studies representing the importance of each node in the network based on certain criteria (Sosa et al. 2021). The node-level network metrics included in the analysis are degree, strength, betweenness, clustering coefficient, eigenvector centrality, harmonic centrality, reach, and laplacian centrality. See Table 3 for a summary of what each of these metrics represents in a network.

**Table 3:**
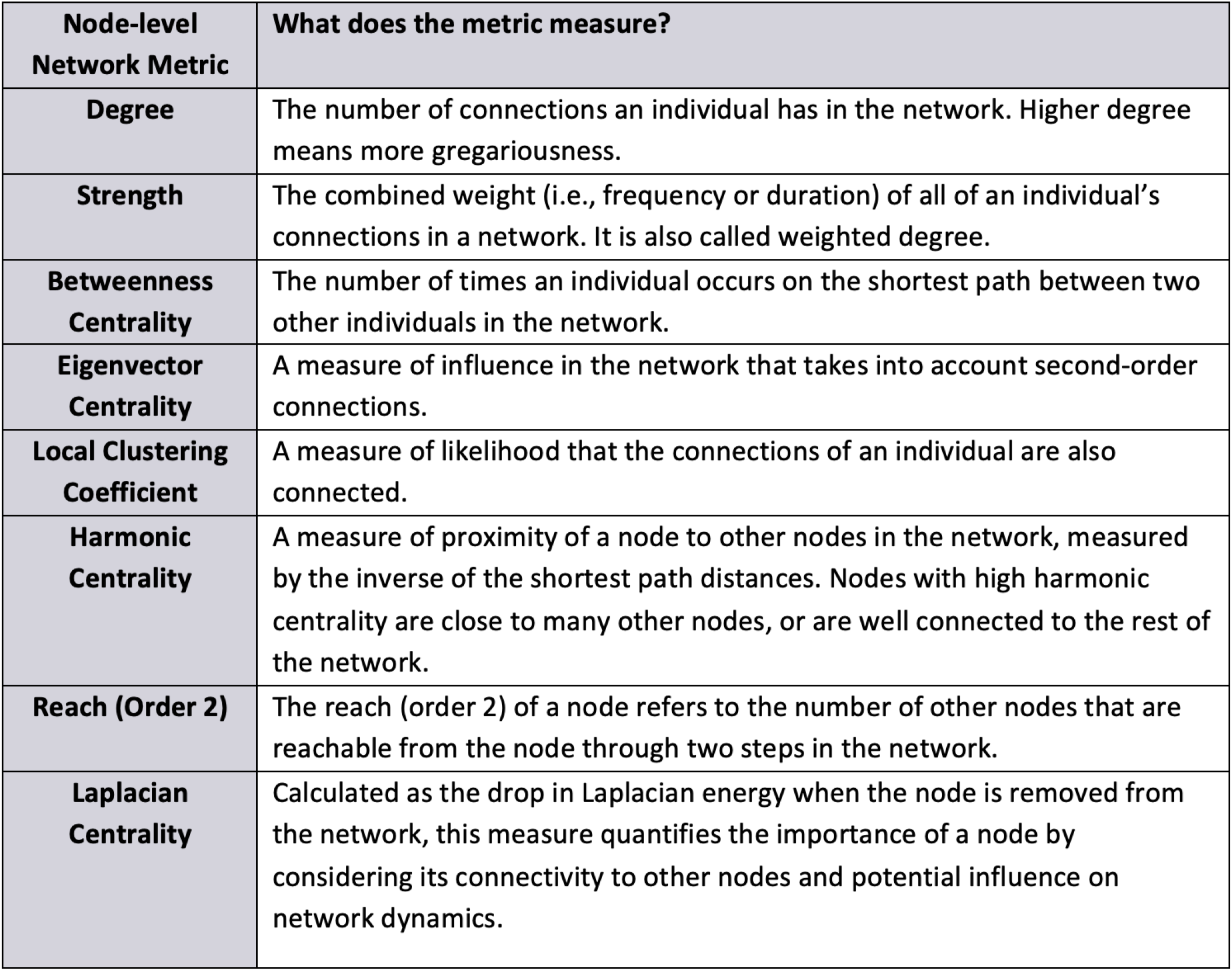
Node-level network metrics used in the analysis of Pronghorn network.

In the fourth step, we examine the correlation coefficient’s behaviour with respect to the node-level network metrics of the observed network and its sub-samples. A significant correlation between the two values shows a strong linear connection and rank preservation among the sampled individuals. The network metrics with a high correlation value provide a more accurate idea of the individual’s position in the social network of the population.

The package function `correlation_analyze()` allows for this analysis on the network obtained from GPS telemetry observations. The function takes in the `igraph` object of the observed network as the first argument, the number of simulations to obtain the mean and standard deviation of correlation coefficient at each subsampling level as the second argument `n_simulations`. The user also specifies the proportions at which sub-sampling should be done via the argument `subsampling_proportion` along with the `network_metrics` to be calculated.

~~~
*R> correlation_pronghorn <-correlation_analyze(pronghorn_network,
     n_simulations = 10,
     subsampling_proportion = c(0.1, 0.3, 0.5, 0.7, 0.9),
    network_metrics_functions_list =
       c(“Degree” = function(net, sub_net)
           igraph::degree(net, v = igraph::V(sub_net)$name),
       “Strength” = function(net, sub_net)
         igraph::strength(net, v = igraph::V(sub_net)$name),
       “Betweenness” = function(net, sub_net)
          igraph::betweenness(net, v = igraph::V(sub_net)$name),
      “Clustering coefficient” = function(net, sub_net)
          igraph::transitivity(net, type = “local”,
            vids = igraph::V(sub_net)$name),
      “Eigenvector centrality” = function(net, sub_net)
          igraph::eigen_centrality(net)$vector[igraph::V(sub_net)$name],
      “Harmonic centrality” = function(net, sub_net)
         igraph::harmonic_centrality(net, vids = igraph::V(sub_net)$name),
      “Reach (order 2)” = function(net, sub_net)
         igraph::ego_size(net, order = 2, nodes = igraph::V(sub_net)$name),
     “Laplacian centrality” = function(net, sub_net)
        centiserve::laplacian(net, vids = igraph::V(sub_net)$name))
        )
R> plot(correlation_pronghorn)*
~~~

The function `correlation_analyze()` returns an object of class `list_correlation_matrices` which contains a list of size equivalent to the number of network metrics specified. Corresponding to each network metric, a matrix is returned that contains the correlation coefficient value at each run of simulation for each subsampling proportion. The returned object can be plotted via the `plot()` function and visualisation is obtained as shown in Figure 9.

**Figure 9.**
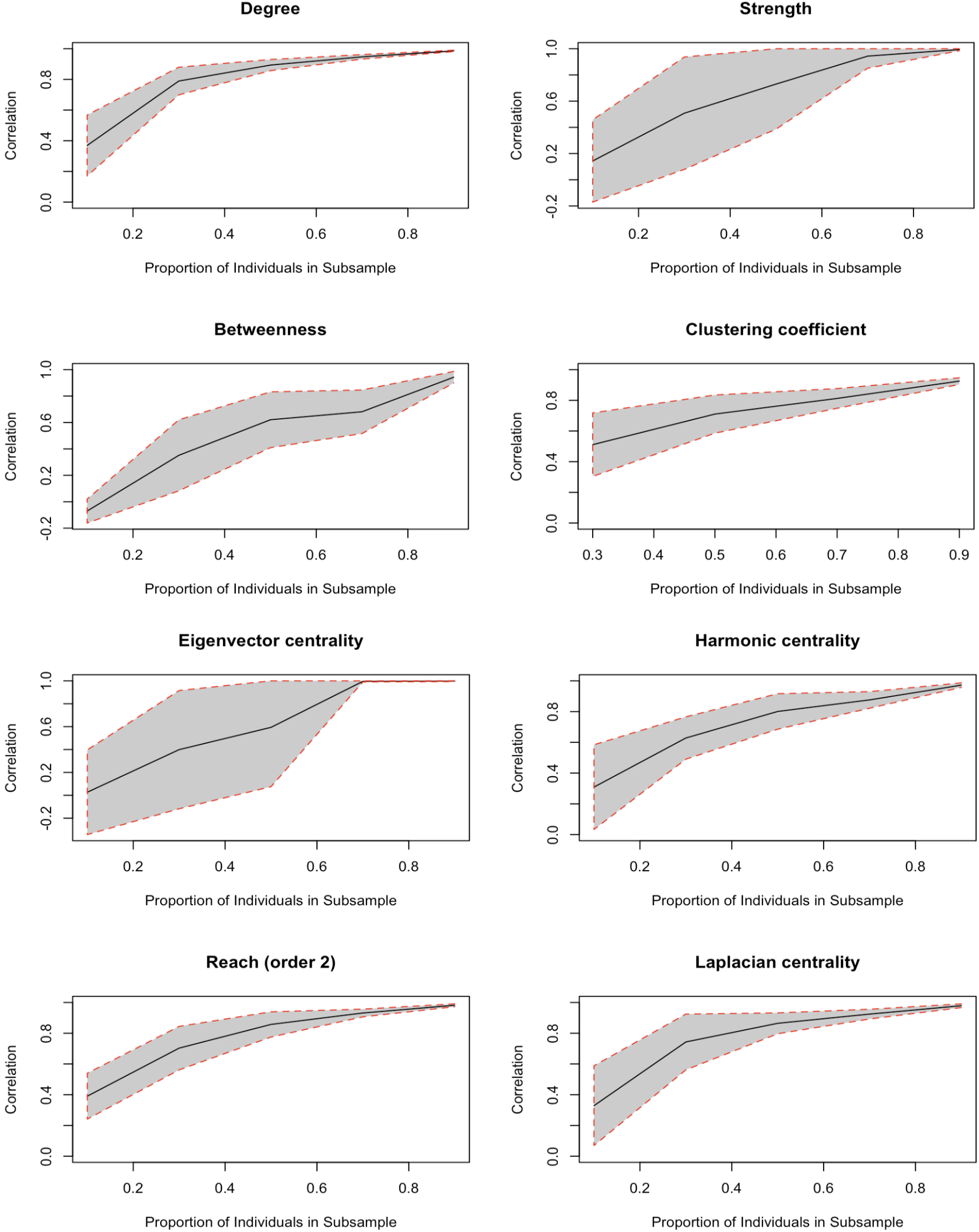
The plots show the correlation of node-level network metrics between the nodes of sub-sampled and observed networks. The black line in the plots indicates the mean correlation coefficient value between the node-level metrics of nodes present in the sub-sampled network. The colored region depicts the standard deviation of the correlation values at each sampling level.

For the pronghorn network, even with a sample size of only 40%, the network measures degree, reach, and laplacian centrality remain well correlated. The correlation values for strength and eigenvector centrality vary greatly as the subsampling level decreases as indicated by the thick band of standard deviation in the value. This indicates that for the pronghorn network, even if 50% of the current sample size was available, the individual rankings for the network metrics degree, reach, and laplacian centrality would have been preserved. As the laplacian centrality of a node represents its influence over the entire network, taking into account both the node’s direct connections and its indirect influence through neighboring nodes, this result implies that the pronghorn network structure remains well preserved when the sample size is lowered to 50% of the current size.

#### Step 4b : Regression analysis between the node-level network metrics of observed and smaller subsamples

Regression provides a quantitative measure of the strength and direction of the relationship between two quantities. We use regression analysis to analyse how the dependence of node-level metrics in sub-sampled networks on the nodes of fully observed network changes when there is a decline in the sampling proportion. The package function `regression_slope_analyze()` allows to perform regression analysis where the values of node-level network metrics from sub-sampled networks are regressed on the node-level network metrics of the same nodes from the observed networks. We show the example of performing regression analysis on the pronghorn network.

~~~
*R> regression_pronghorn <-regression_slope_analyze(pronghorn_network,
           n_simulations = 10,
           subsampling_proportion = c(0.1, 0.3, 0.5, 0.7, 0.9),
          network_metrics_functions_list =
    c(“Degree” = function(net, sub_net)
          igraph::degree(net, v = igraph::V(sub_net)$name),
    “Strength” = function(net, sub_net)
          igraph::strength(net, v = igraph::V(sub_net)$name),
    “Betweenness” = function(net, sub_net)
          igraph::betweenness(net, v = igraph::V(sub_net)$name),
    “Clustering coefficient” = function(net, sub_net)
         igraph::transitivity(net, type = “local”,
            vids = igraph::V(sub_net)$name),
    “Eigenvector centrality” = function(net, sub_net)
           igraph::eigen_centrality(net)$vector[igraph::V(sub_net)$name],
    “Harmonic centrality” = function(net, sub_net)
         igraph::harmonic_centrality(net, vids = igraph::V(sub_net)$name),
     “Reach (order 2)” = function(net, sub_net)
         igraph::ego_size(net, order = 2, nodes = igraph::V(sub_net)$name),
     “Laplacian centrality” = function(net, sub_net)
         centiserve::laplacian(net, vids = igraph::V(sub_net)$name)))
R> plot(regression_pronghorn)*
~~~

The function `regression_slope_analyze()` returns an object of class `list_regression_matrices` which can be passed to the `plot()` function to obtain a visualisation as shown in Figure 10. For the pronghorn network, the slope of regression declines almost linearly with a decrease in the sampling proportion for the network metrics degree. It follows an almost linear pattern for strength, harmonic centrality, reach, and laplacian centrality. This shows that for the sub-samples of the available sample of pronghorn, the variation in the rank orders for the individuals in terms of the network metrics degree, harmonic centrality, reach, and laplacian centrality can be explained well, however, the strength of this relationship declines as the sample size is lowered. The network metric clustering coefficient has a regression slope of 1 when as low as 30% of the pronghorn observed sample is sub-sampled. Therefore, rankings based on the clustering coefficient of smaller sub-samples reflect the true rankings of the observed sample. As the observed sample represents just a proportion of the population, the network metrics that do not preserve the rankings should not be used to make inferences on the population’s network structure.

**Figure 10.**
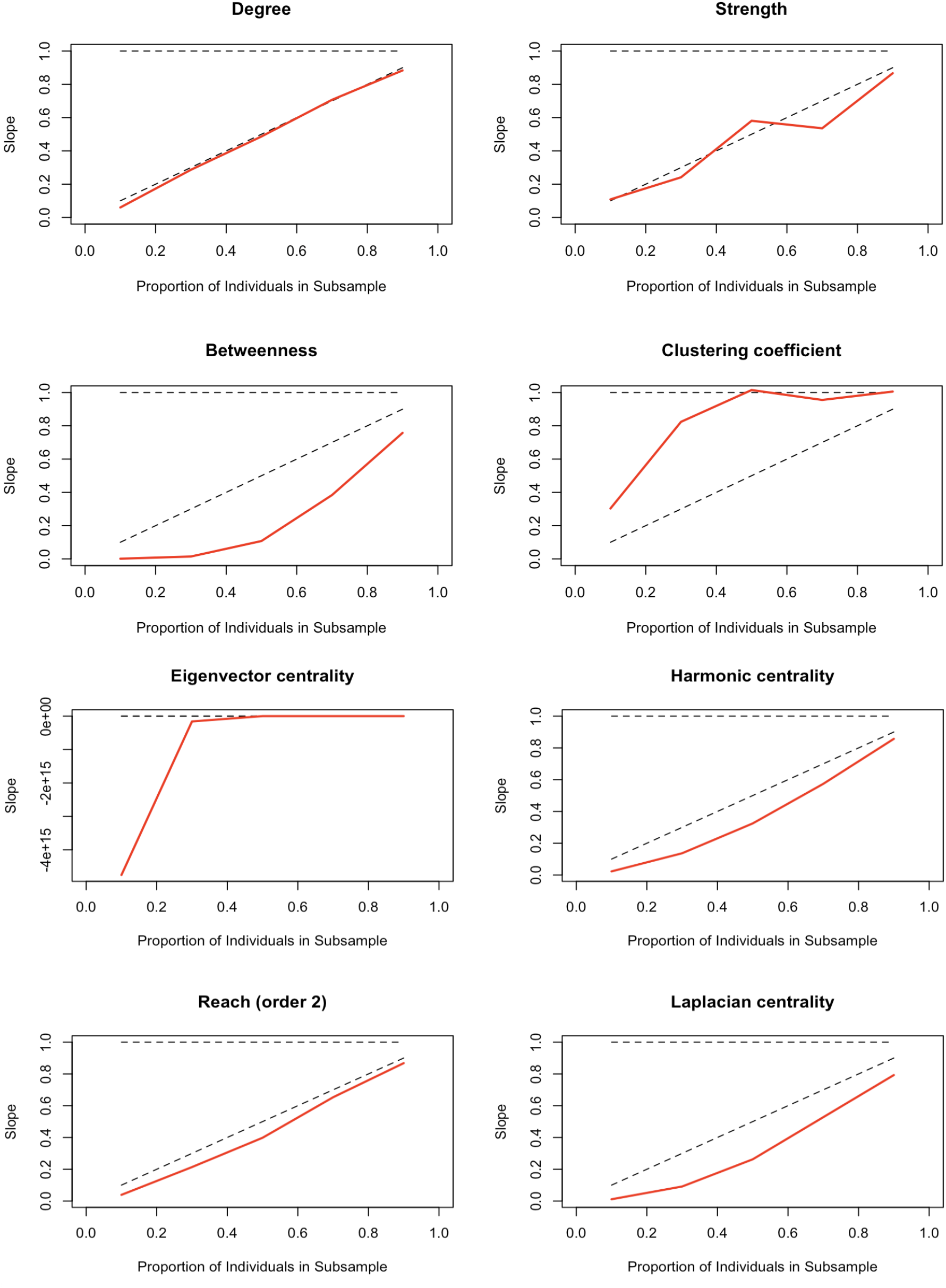
Regression analysis of the node-level network metrics between the sub-sampled and observed network. In each plot, the x-axis denotes the proportion of nodes in the sub-sample and the y-axis shows the corresponding value of the slope of regression calculated by regressing these metrics of sub-sampled nodes on the observed network nodes.

#### Step 5 : Confidence intervals for node-level network metrics

As with global network metrics, confidence intervals play an important role in enhancing network analysis inferences for node-level network metrics. For each individual in the sample, point estimates provide a snapshot of their position in the network which might change depending on the choice of other individuals in the network. It is important to assess the extent of this change when dealing with a sample from the population. Confidence intervals provide a range of values within which the metric value of the node is likely to fall. The final step in the five-step workflow is to obtain confidence intervals around the point estimates of the node-level network metrics. The package function `node_level_CI()` allows the generation of confidence intervals for each node’s observed network metric value.

~~~
*R> pronghorn_node_level_CI <-node_level_CI(pronghorn_network,
           n_versions = 100,
           network_metrics_functions_list =
                 c(“Degree” = igraph::degree,
                 “Strength” = igraph::strength,
                 “Betweenness” = igraph::betweenness,
                 “Clustering coefficient” = function(x){
                       trans <-igraph::transitivity(x,
                       type = “local”, vids = igraph::V(x),
                       isolates = “zero”);
                       names(trans) <-igraph::V(x)$name;
                       return(trans)},
                 “Eigenvector centrality” = function(x)
                     igraph::eigen_centrality(x)$vector,
                 “Harmonic centrality” =
                         igraph::harmonic_centrality,
                 “Reach (order 2)” = function(x){
                    reach <-igraph::ego_size(x, order = 2)
                   names(reach) <-igraph::V(x)$name
                   return(reach)
                   },
           “Laplacian centrality” = centiserve::laplacian),
    n_cores = 1, CI_size = 0.95)
R> plot(pronghorn_node_level_CI)*
~~~

The package function `node_level_CI()` takes in the observed network in the form of an `igraph` object. The user is asked to specify the number of bootstrapped versions to be considered for obtaining the confidence intervals with a default value of 100. The user can also specify the level of confidence through `CI_size` argument. The default value is 0.95 which generates 95% confidence intervals. The function returns a list which is an object of class `list_node_level_CI`. Passing this list to the plot function generates visualisation as depicted in figure 11.

**Figure 11.**
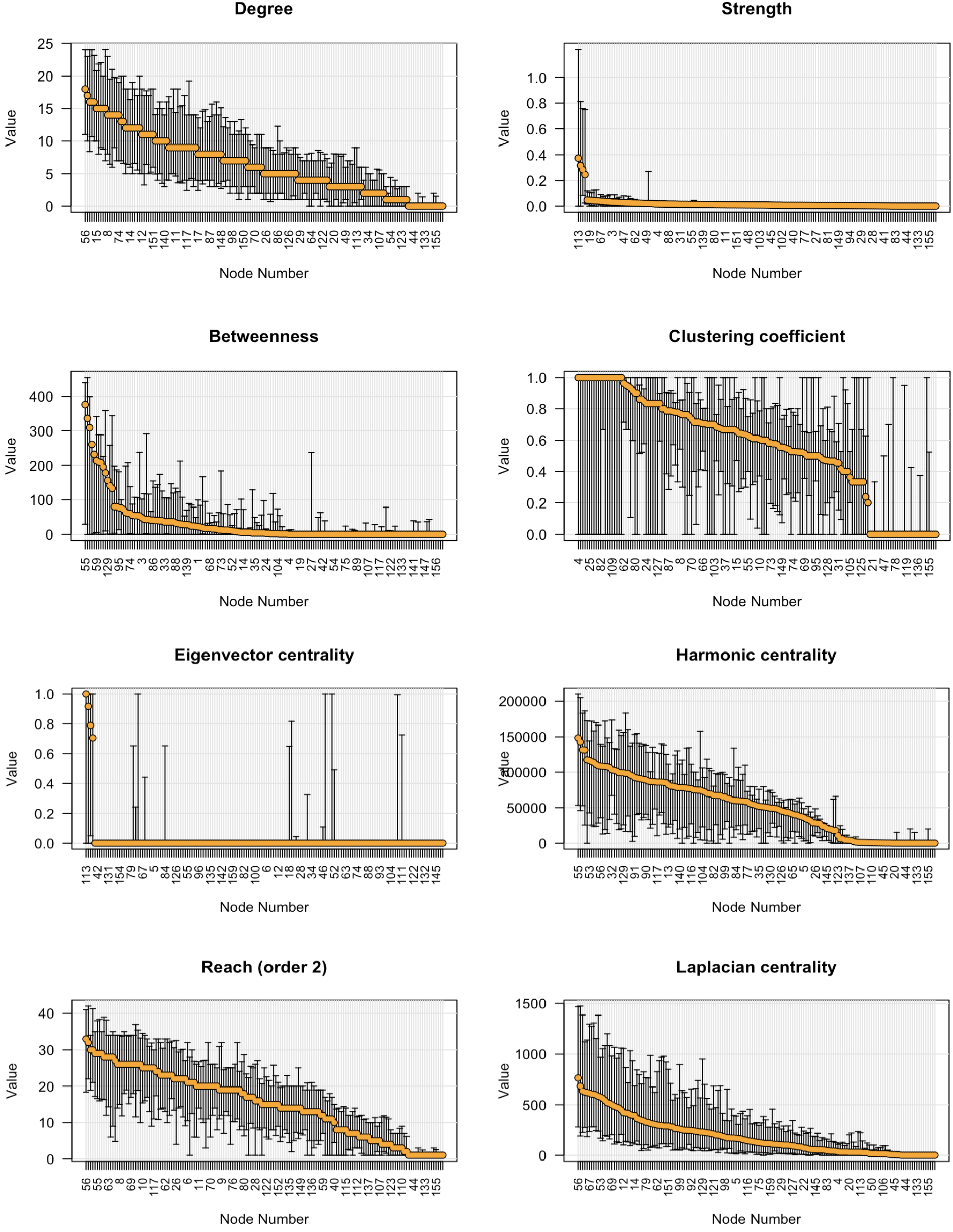
Node-level metrics for the pronghorn network for the network metrics degree, strength, betweenness, clustering coefficient, eigenvector centrality, harmonic centrality, reach, and laplacian centrality with associated 95% confidence intervals. To facilitate readability, the nodes are sorted in decreasing order by observed metric.

For the pronghorn network, each of the plot for a network metric depicts an interesting picture. The amount of uncertainity in the observed values of degree for each node is approximately equal. A lot of the nodes having high clustering coefficient values have the tendency to achieve very low values and vice versa depending on the choice of other individuals in the sample. For the network metric eignevector centrality, most of the nodes have an observed value of zero, however we can point out the few nodes from the sample that have a tendency to have higher values for this centrality. Plot for harmonic centrality indicates that some of the nodes have a much higher tendency to act as bridges connecting different parts of the network than what is observed from the given sample.

### Conclusions from pronghorn GPS telemetry data

A summary of the workflow is provided in figure 12. Performing this five-step workflow on the pronghorn network obtained from the GPS telemetry observations has revealed interesting insights about the nature of the observed sample. Along with providing a comprehensive understanding of the network structure, it has helped us to familiarise with the patterns, performance with different social network metrics, and potential issues in the data before proceeding with a more advanced form of social network analysis.

**Figure 12.**
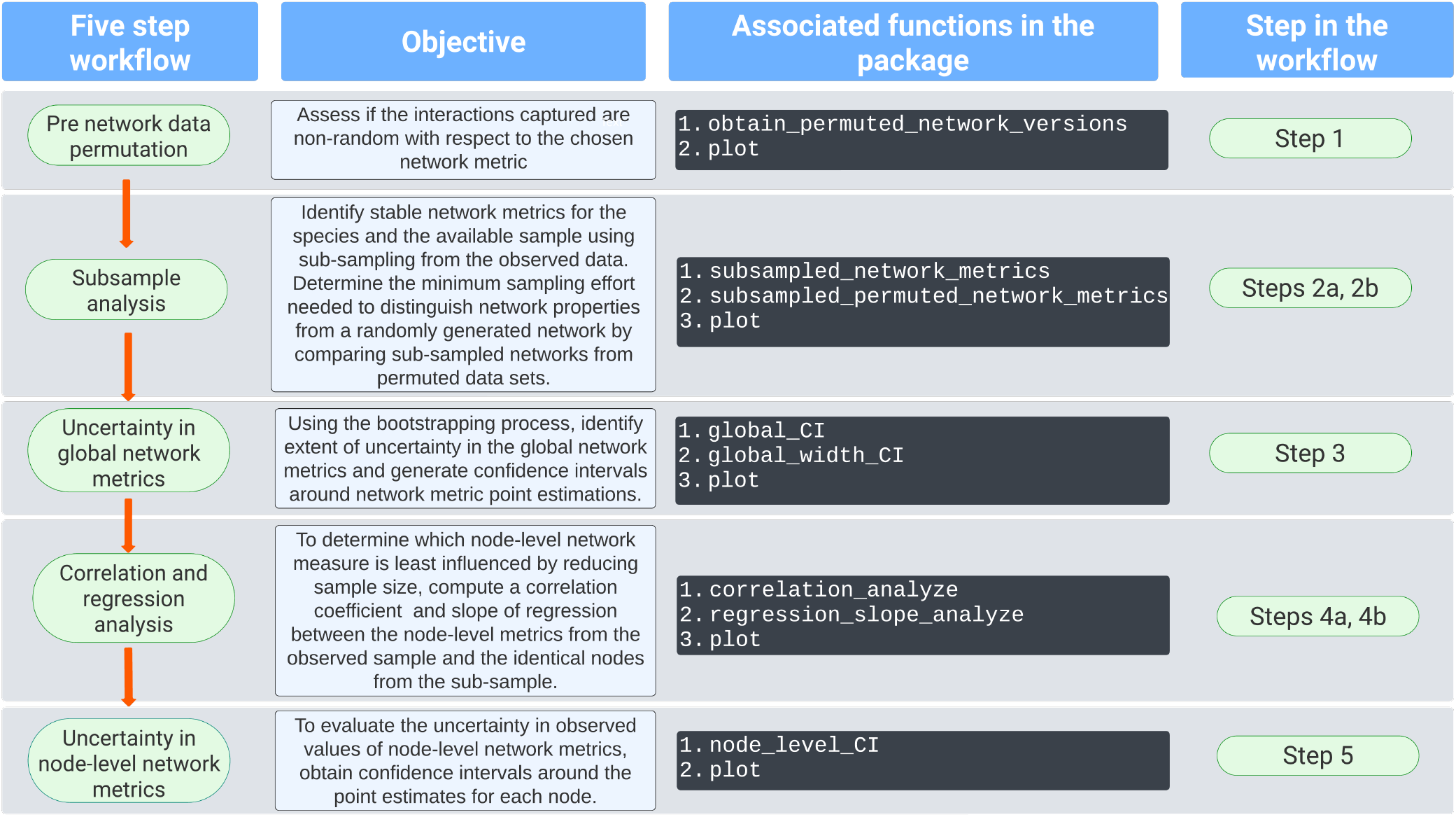
Summary of the five-step workflow

We assessed seven global network metrics and eight node-level network metrics for their compatibility with the raw GPS telemetry observations collected for pronghorn. The first step of the workflow highlighted the network metrics which could be an ideal choice to be used for further social network analysis on the observed pronghorn data. For example, if a researcher were considering using modularity as a network metric to test certain hypotheses, the first step of the workflow revealed that it might not be a good idea. The observed value of modularity lied within the distribution of null values that indicated that the pronghorn network generated from the sampled observations does not explicitly capture this aspect and is in fact similar to a random network with regards to modularity. This is an important result as if a researcher was interested in comparing the network structure of pronghorn in winter and summer months and no difference in modularity was observed. The researcher might conclude that there is no difference in the two networks in terms of modularity whereas the actual reason could be that the data collected does not capture this aspect very well in the first place.

The second and third steps of the workflow revealed insights about the extent of bias in the chosen network metrics and the nature of conclusions that could be made using those. The network metrics edge density, transitivity, and assortativity degree remained unbiased with lowering sample sizes and obtaining confidence intervals around the observed values for these metrics provided a range of values within which the population parameter may lie. The fourth and fifth steps of the workflow assessed the uncertainty in the observed values of node-level network metrics. Degree, reach and laplacian centrality of smaller sub-networks were well correlated with the observed network, indicating that node-wise ranks were well preserved for these network metrics. Having confidence intervals around the observed values of node-level network metrics helped point out the network metrics that tend to achieve much greater (or lower) observed values, depending on the choice of other individuals in the sample. For example, if we need to choose individuals to be vaccinated in cases of disease spread, the individuals from the sample having higher upper confidence interval values for harmonic centrality should be chosen as the individuals with higher harmonic centrality in the sub-network are likely to have highest harmonic centrality in the full population as indicated by high correlation coefficients.

### Computational limitations

Computing pre-network data stream permutations is computationally demanding and could take long depending on the number of individuals monitored, observation frequency and duration of observations. The users are advised to use multiple cores to allow for parallel processing and use remote servers whenever available.

## 4. Discussion

GPS telemetry data is becoming increasingly common for extracting and analyzing animal social networks, providing valuable insights into the social and spatial behavior of animal communities. By leveraging the latest tools available for animal social network analysis, researchers can uncover nuanced insights into how animals interact socially and navigate their environments. In this endeavour, the aniSNA serves as a crucial but thus far missing tool that is needed before using many existing tools for SNA on animal datasets. It allows the user to construct and analyze network structures even when working with a subset of the population. This means that data previously considered insufficient for social network analysis can now be judiciously utilized, expanding the scope of research possibilities. The package includes sample data from GPS telemetry observations of elk monitored in 2010, which users can use to explore the methods described in this paper for themselves.

The package aniSNA will continue to grow as further advancements in statistical tools to analyse and obtain inferences from network data take place. The current and subsequent versions of aniSNA enable the researchers to derive robust statistical insights from the networks obtained from GPS telemetry data. Animal ecologists gain the capacity to compute an array of social network metrics, offering insights at both population-wide patterns and individual behaviors. It enables users to confidently evaluate the reliability of these metrics and leverage them for subsequent analyses. For instance, researchers can investigate the intricacies of social network variation within and between populations, revealing how social structures differ in various ecological environments. Additionally, by linking individual sociality metrics to life history traits, such as reproductive success or survival rates, they can uncover the profound implications of social dynamics on key aspects of animal behavior and ecology. Through these capabilities, the package serves as a powerful tool for unraveling the complexities of animal social systems and their ecological implications, paving the way for more informed conservation and management strategies.

## Acknowledgments

This publication has emanated from research conducted with the financial support of Science Foundation Ireland under Grant number 18/CRT/6049. For the purpose of Open Access, the author has applied a CC BY public copyright licence to any Author Accepted Manuscript version arising from this submission.

## Data availability

Should the manuscript be published, the data will be made publicly available to allow analyses to be fully reproducible. GPS locations, however, will be randomly jittered or censored due to data embargo related to the single research projects.

## Author contributions

Prabhleen Kaur and Michael Salter-Townshend conceived the ideas and designed the methodology with the help of Simone Ciuti. Prabhleen Kaur analysed the data and wrote the manuscript, edited by Michael Salter-Townshend and Simone Ciuti. Adele K. Reinking and Jeffrey L. Beck provided the data and contributed to the revision of the manuscript. All authors contributed critically to the drafts and gave final approval for publication.

